# Behavioural flexibility and its drivers in semi-urban vervet monkeys

**DOI:** 10.1101/2025.06.06.658231

**Authors:** Paige Barnes, Benjamin Robira, Stephanie Mercier, Sofia Forss

## Abstract

Phylogenetic comparisons suggest that behavioural flexibility facilitates success in urban environments. It remains less clear whether urbanization fosters cognitive skills that require flexibility, or whether successful individuals in urban environments simply apply pre-evolved skills to solve new problems. To investigate whether variation in anthropogenic experience drives behavioural flexibility required to solve a new technical problem, we presented 42 semi-urban vervet monkeys (*Chlorocebus pygerythrus*) with a three-phase foraging experiment. For this, we measured each monkey’s tendency to “raid” human structures in their search for food to assess whether their anthropogenic experience explained individual variation in flexibility and problem-solving skills. We then used a probabilistic model to describe the series of monkeys’ actions when attempting to solve the foraging experiment. This allowed us to quantify three traits encompassing individuals’ behavioural flexibility: switch tendency between solutions, innovativeness, and learning sensitivity. We found that individuals’ switch tendency and innovativeness were not explained by anthropogenic experience and that the three traits were unrelated to each other. Moreover, neither switch tendency, nor innovativeness predicted the monkeys’ human food consumption in this habitat. Thus, our study suggests that behavioural flexibility is not driven by anthropogenic foraging experience. Contrasting with the hypothesis that urbanization selects for behavioural flexibility, our findings instead imply that vervet monkeys already had the sufficient behavioural flexibility and cognitive capacities to successfully exploit the urban habitat.

## 1. Introduction

When animals encounter a novel problem, they can either persistently pursue a previously used behaviour (i.e., *stereotypy*) or exhibit behavioural flexibility. Behavioural flexibility may involve either switching to another known and effective behaviour (i.e., *switch tendency*; (1)) or modifying the repertoire resulting in a new behaviour (i.e., *innovativeness*; (2,3), but see (4)). Under changing conditions, only switch tendency and innovativeness are adaptive (3,5,6), while stereotypy is not. Effective switching or innovating requires animals to act or build upon their repertoire of useful behaviours, highlighting the role of remembering past actions (i.e., *learning sensitivity*; (7,8)).

On an evolutionary timescale, urban environments represent novel environments. Phylogenetic comparisons show that behaviourally flexible species are more able to cope with anthropogenic challenges, making them overall more likely to thrive in urban habitats (9–12). However, given the evolutionary recency of urbanization, there is limited evidence of adaptation to these environments. In fact, some studies (e.g., (13)), find little to no signs of behavioural adaptive selection. Therefore, while flexibility may support persistence in urban environments, it is unclear whether urban life selects for cognitive traits or merely draws upon existing behavioural repertoires. While interspecific comparisons demonstrate that flexible species are more likely to persist in urban landscapes, it remains unclear whether urban experience in turn shapes the cognitive traits of individuals.

Within-species comparisons along the urban-rural gradient have used experimental foraging tasks to specifically quantify behavioural flexibility and test its evolutionary relevance in urban environments, but these have revealed contrasting results. Urban individuals sometimes appeared more innovative (field mice, *Apodemus agrarius*: (14); bullfinches, *Loxigilla barbadensis*: (15); great tits, *Parus major*: (16)), while other findings report equal or lower innovativeness compared to their rural counterparts (foxes, *Vulpes vulpes*: (17); grey squirrels, *Sciurus carolinensis*: (18); spotted hyenas, *Crocuta crocuta*: (19); multiple bird species: (20,21)). Such contrasting results have, however, to be considered cautiously as absence of evident differences is not evidence of absence of differences. Indeed, behaviours may be latent and only expressed when required (22,23). Apparent urban and rural differences in cognitive performance may reflect variation in task engagement or motivation, rather than differences in cognitive capacities (24–26). As such, these inconsistent findings from urban-rural comparisons highlight the need to examine how individual variation in anthropogenic experience relates to behavioural flexibility.

Whilst we know that, within populations, individuals’ social and physical environmental experiences, such as predation pressures, food availability, contact with humans, and enrichment, can impact cognitive phenotypes (27–29), less is known about how anthropogenic experiences, such as exploiting urban food sources, affect individuals’ cognitive skills. If urban environments promote behavioural flexibility, we expect individuals with greater exposure to anthropogenic foraging opportunities, often requiring manipulation of manmade structures (30), to show enhanced problem-solving abilities. On the contrary, if urban environments do not promote cognitive skills necessary to flexibly solve novel problems, this would suggest that the species’ pre-existing flexibility and latent cognitive capacity is sufficient for survival, and consequently, individual variation in cognitive abilities stem from other drivers.

In this study, we investigated whether behavioural flexibility emerges from exposition to urban foraging experiences in a semi-urban vervet monkey (*Chlorocebus pygerythrus*) population living in the Simbithi Eco-Estate in South Africa. Vervet monkeys are known to be habitat and food generalists and able to adopt flexible foraging strategies (31–34). Accordingly, part of their successful coping with anthropogenic disturbance could be associated with the species’ ability to profit from human food resources (35,36). Consequently, we investigated three questions: (i) Does anthropogenic exploitative foraging (hereafter referred to as “raiding tendency”) in this species transfer to individual variation in problem-solving skills (requiring behavioural flexibility)? (ii) Do flexible traits, such as “switch tendency”, “innovativeness” and “learning sensitivity,” co-vary to form a behavioural syndrome? (iii) Does individual variation in behavioural flexibility relate to (a) successful solving of a technical challenge, and (b) increased human food consumption (as a proxy for successful exploitation of the anthropogenic habitat)?

Given that switch tendency, innovativeness, and learning sensitivity are all aspects of behavioural flexibility and therefore argued to be advantageous in urbanized environments, these traits may also be driving the monkeys’ success in this habitat. Consequently, if co-variations occur between the three traits, they imply that selection may have favoured a set of adaptive traits forming a so called “syndrome” (37,38), as observed for other adaptive trait clusters like aggression-boldness (39). Selection in urban habitats may favour such an adaptive behavioural flexibility syndrome if the three traits jointly enhance foraging success and reproductive output. Alternatively, urban pressures may favour only specific flexibility components at the cost of another, such as high innovativeness and learning sensitivity, but persistence instead of high switch tendency, resulting in trade-offs between traits (40).

Finally, should behavioural flexibility be key to individual success in urbanized habitats, and already undergone positive selection, we expect that (i) switch tendency and innovativeness are associated with an individual’s raiding tendency, (ii) this population of monkeys to show a syndrome representing high switchers, innovative, and sensitive learners, and (iii) high-switching and innovative individuals should (a) perform better in the foraging experiment and (b) consume more human food.

To answer these questions, we presented 42 individually identified monkeys with a three-phase foraging experiment (puzzle box) of increasing technicality. We used conditional probabilistic modelling to consider the whole temporally-ordinated series of each monkey’s interactions with the puzzle box. This approach enabled us to evaluate decision choices by characterizing three distinct measures of each individual’s behavioural flexibility in the experiment: switch tendency (shifting between solutions), innovativeness (opening different options) and learning sensitivity (repeating successful techniques). Finally, we also measured each monkey’s success at accessing the food reward.

## 2. Methods

### 2.1 Study population

We studied two troops of semi-urban vervet monkeys (N = 42, including 13 adult females, four adult males, and 25 juveniles) ranging in Simbithi Eco-Estate in Ballito, KwaZulu Natal province, South Africa. Simbithi Eco-Estate is a residential, gated community with endemic vegetation on forested trails creating a mosaic of forest patches, human-associated vegetation, houses, and impervious surfaces (e.g., roads), all of which are used by the vervet monkeys (41). Both troops were habituated to human observers and behaved naturally in the presence of researchers.

### 2.2 Foraging experiment

#### 2.2.1 Experimental set-up

The foraging experiment featured a puzzle box with a two-option design (33). This transparent plastic box had the back, bottom, and adjacent sides painted in blue (dimensions: 13 × 10.5 × 7.9 cm) and could be opened in two different ways to gain access to a food reward (i.e., a peanut) inside: by “lifting” a lid or “pulling” the draw container (figure 1, photos in figure S1).

**Figure 1.**
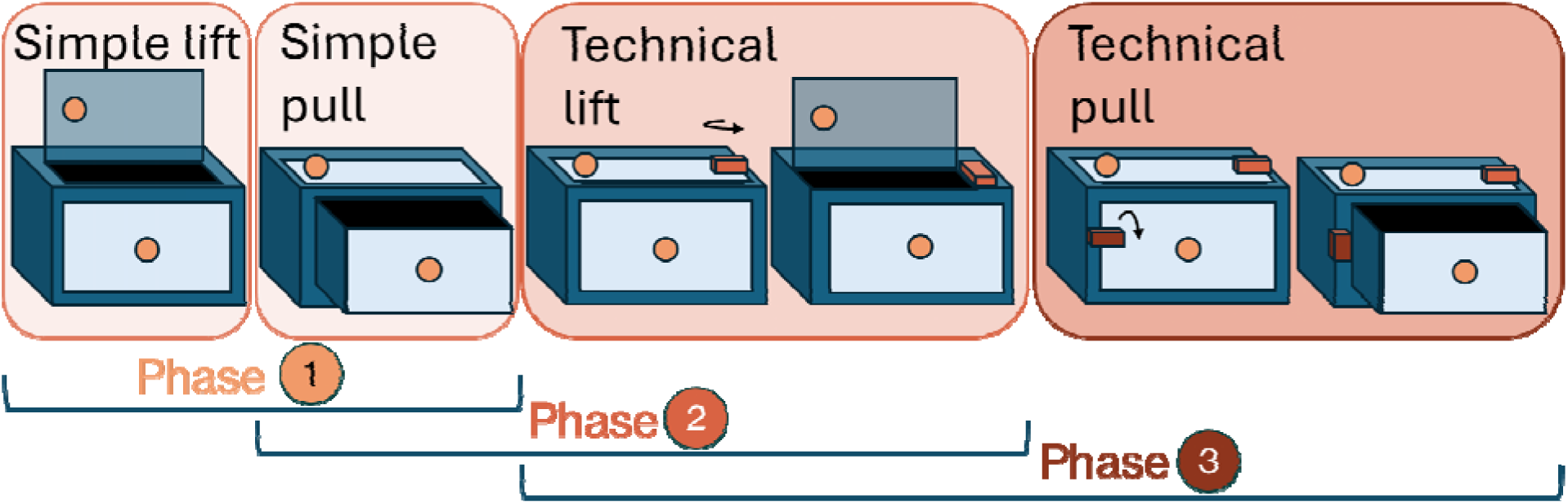
Visualization of the two options for each of the three phases of the experiment. The colours of the phase and box illustrations correspond to the technicality of the phase due to added opening obstacles along the experimentation, with the darker the graphic illustration, the more technical. Phase 1: the monkey can simply either lift (“simple lift” solution) or pull (“simple pull” solution); Phase 2: the monkey can either use the simple pull or unlock a small block at the side of the lid and lift (“technical lift solution); Phase 3: the monkey can either use the technical lift or technical pull solution (incorporating first unlocking a block at the side of the lid and/or at the side of the drawer).

Experimental data were collected from September 2023 to January 2024. During this time, we opportunistically presented the monkeys with the three-phase puzzle box foraging experiment. Experiments were conducted at any time throughout the day from sunrise to sunset. To avoid multiple individuals interacting with the box and inter-individual conflicts, we specifically seized opportunities to target one individual and perform the experiment when a limited number of individuals were present. To set up the experiment, we anchored the puzzle box to the ground using camping hooks and placed the box, so it was visible to the target monkey. We changed the direction of the box when moving location between trials to avoid possible constant directional bias on side manipulation caused by the potential influence of an observer’s presence at a given side. All experiments were videotaped (see 2.2.2.4).

Since these monkeys were naïve to this experimental setup, we first implemented a pre-exposure phase (familiarization phase) to minimise any potential neophobia towards the box, which could impact the likelihood of interacting with it (42). During the familiarization phase, there was no obstacle to accessing the food reward as we removed the top lid and pull drawer of the box and baited it with a peanut on top of the box so that the monkeys could form a positive association to the puzzle box. We ended the familiarization phase once every individual in both troops retrieved a peanut, and all troop members were determined to have been within eye line of at least one retrieval. The experiment thereafter comprised three phases, each implying a different level of technicality (figure 1). Subsequently, in a few situations, we continued to bait the top of the box with an additional peanut to attract the attention to the puzzle box when an individual showed low motivation to participate.

A trial began when a monkey touched the puzzle box and ended when it moved out of arm’s length for over a minute (all phases), or it reached a one-minute time limit while within arm’s length of the box, interacting or not (phases 2 and 3). Groups were limited to testing up to twice a week on non-consecutive days to avoid overfeeding and to minimize potential behavioural impacts due to the experiment. An individual was tested a maximum five times a day (mean ± sd: 1.99 ± 1.12; with testing spanning over 97.10 ± 41.71 days) and the time interval between trials was of a minimum of one minute if unsuccessful, but null if successful (a new peanut was immediately reinserted in the box). We considered a trial as “successful” when the participant successfully retrieved the peanut, or otherwise as an “attempt” if not.

Following an ethogram, (electronic supplementary material, table S1), we used video records to thoroughly identify whether unsuccessful access to the peanut could be considered as attempts nonetheless (i.e., the individual tried to open the box but did not manage to, in contrast to when a monkey ignored or only superficially explored the box). If the interaction with the box was interrupted by a conspecific, we considered this trial as “aborted” and it was not further included in the analyses. All individuals were free to engage in each trial or not (electronic supplementary material, table S2). This means that, for each trial, we considered four possible outcomes: successfully lifting, successfully pulling, unsuccessfully lifting, and unsuccessfully pulling. Only unsuccessful lift and pull were non-exclusive, meaning that if an individual attempted a solution (i.e. was unsuccessful) at first but was later successful, we only considered the successful event, and the trial could be labelled with the outcome “successful lifting.” However, if the individual attempted both lifting and pulling, the trial would be labelled “unsuccessful lifting and pulling.” Attempts were, however, only counted as “one” (i.e., we did not consider whether the individual attempted one or several times an opening option within a trial), for the sake of interpretability.

Each phase consisted of several trials (N_phase1_ = 444 trials and 41 monkeys (one monkey never engaged in the experiment), N_phase2_ = 174 and 27 monkeys, N_phase3_ = 155 and 25 monkeys). While all individuals could participate in phase 1, phases 2 and 3 were constrained to individuals who passed a “phase success” criterion each time. To validate our chosen phase success criterion, the influence of the number of trials per phase on our estimations can be found in the electronic supplementary material (figures S4 – S6).

##### 2.2.2.1 Phase 1 – Simple lift and simple pull

In phase 1, the lid and the drawer, absent in the familiarization phase, were added, such that the peanut could be retrieved by the simple solutions of either “pulling” the drawer or “lifting” the lid (figure 1). We considered that a monkey passed phase 1 when it achieved 10 trial successes (consisting of either lifting or pulling, or a mixture of both solutions), as this indicated that the monkey could adequately perform at least one simple solution, and this number provided enough trials to robustly examine switch tendencies. Monkeys were free to try the experiment as many times as they wanted but only passed phase 1 if they reached at least 10 successes.

##### 2.2.2.2 Phase 2 – Technical lift and simple pull

Individuals overall preferred the lift option in phase 1 (83%). In phase 2, we increased the technicality of the “lifting” access by adding a twistable lock (a small white block: 2 x 1 x 1 cm) screwed onto the top of the box blocking the lid (figure 1). When the block was turned in either direction, the lift solution could be “unlocked”, and the lid could be lifted as in phase 1. Because the monkeys had to use fine motoric finger movements or employ teeth to twist the lock to retrieve the peanut, this was considered a “technical” solution compared to simply lifting and pulling the lid. If the monkeys managed to retrieve the reward within a minute, trials were counted as a success, otherwise as an attempt (if the monkey interacted with the box). Individuals who completed a minimum of six trials moved to phase 3.

##### 2.2.2.3 Phase 3 – Technical lift and technical pull

In phase 3, we maintained the “technical” lift solution from phase 2, and increased the difficulty of the pull solution in the same way as the lift solution (figure 1), introducing an equivalent lock on the original pull side. Success and attempts were defined in the same way as phase 2, and phase 3 also ended when the monkeys reached six trials.

##### 2.2.2.4. Video coding

All experimental trials were recorded using a Canon camera HDR-CX200 mounted to a tripod, placed on average about 5 m from the box. Individuals’ identities within the eyeline of the box, and distances to the box (i.e., within 2, 5, 10, or more than 20 m) were recorded vocally during the filming of the experiment. Trial outcomes were subsequently extracted (table S1).

### 2.3 Behavioural observations

We used long-term observational data, where the monkeys were monitored daily from Monday to Friday for up to eight hours between sunrise and sunset, cumulating in 3137 hours of observation time between July 2023 and September 2024. Observers recorded the monkeys’ social and feeding behaviours (electronic supplementary material, table S3) following *ad-libitum* records of specific events (e.g., opportunistic feedings on anthropogenic food resources, referred to as “raiding” behaviour) using handheld phones (Blackview BV4900) and CyberTracker version 3.531. Each observer that contributed to this long-term data set passed both an individual identification and an inter-observer reliability test which had to be successfully completed prior to collecting data with at least 80% of agreement reached, estimated with a Cohen’s Kappa test (electronic supplementary material, table S4 and section S5 for further details).

The tendency to “raid” anthropogenic structures was defined as each time a monkey entered a building or interacted with a bin. We hypothesized that such instances could accumulate the monkeys’ experiences with anthropogenic artefacts, and thereby promote technical problem-solving skills, useful to innovatively solve a manmade puzzle box. Using *ad-libitum* data, we quantified “raiding tendency” as the number of times individual monkeys foraged on trash or entered a house divided by the total amount of follow hours of that troop when that individual was present (range: 3 to 72, mean = 27.8 events per monkey). In addition, we estimated the human food consumption, or the rate at which each individual fed on human food sources by calculating the number of recorded instances an individual was seen to successfully obtain/consume food in the *ad-libitum* data, divided by the corresponding number of observation hours.

### 2.4 Ethical approvals

This project has been approved by the Animal Research Ethics Committee of the South African University of KwaZulu-Natal (protocol reference number: T20220164), as well as supported by the Environmental Board of Simbithi Eco-Estate in KwaZulu Natal province, South Africa.

### 2.5 Statistical analyses

All data processing and analyses were performed with *R* software (v4.4.1, (43)).

#### 2.5.1 Quantifying individuals’ switch tendency, innovativeness, and learning sensitivity

We distinguished “simple” and “technical” challenges (and thus, considered two sets of switching and innovative parameters) to disentangle the effects on the cognitive skills when options vary in cognitive and/or motoric demand. We used probabilistic modelling to characterize five traits for each individual: simple/technical switch tendency, simple/technical innovativeness, and learning sensitivity (table 1). These scores were calculated from using specific decision trees (figure 2 & figure S2) based on the conditions of each trial and updated throughout the entire sequence of trials.

**Figure 2.**
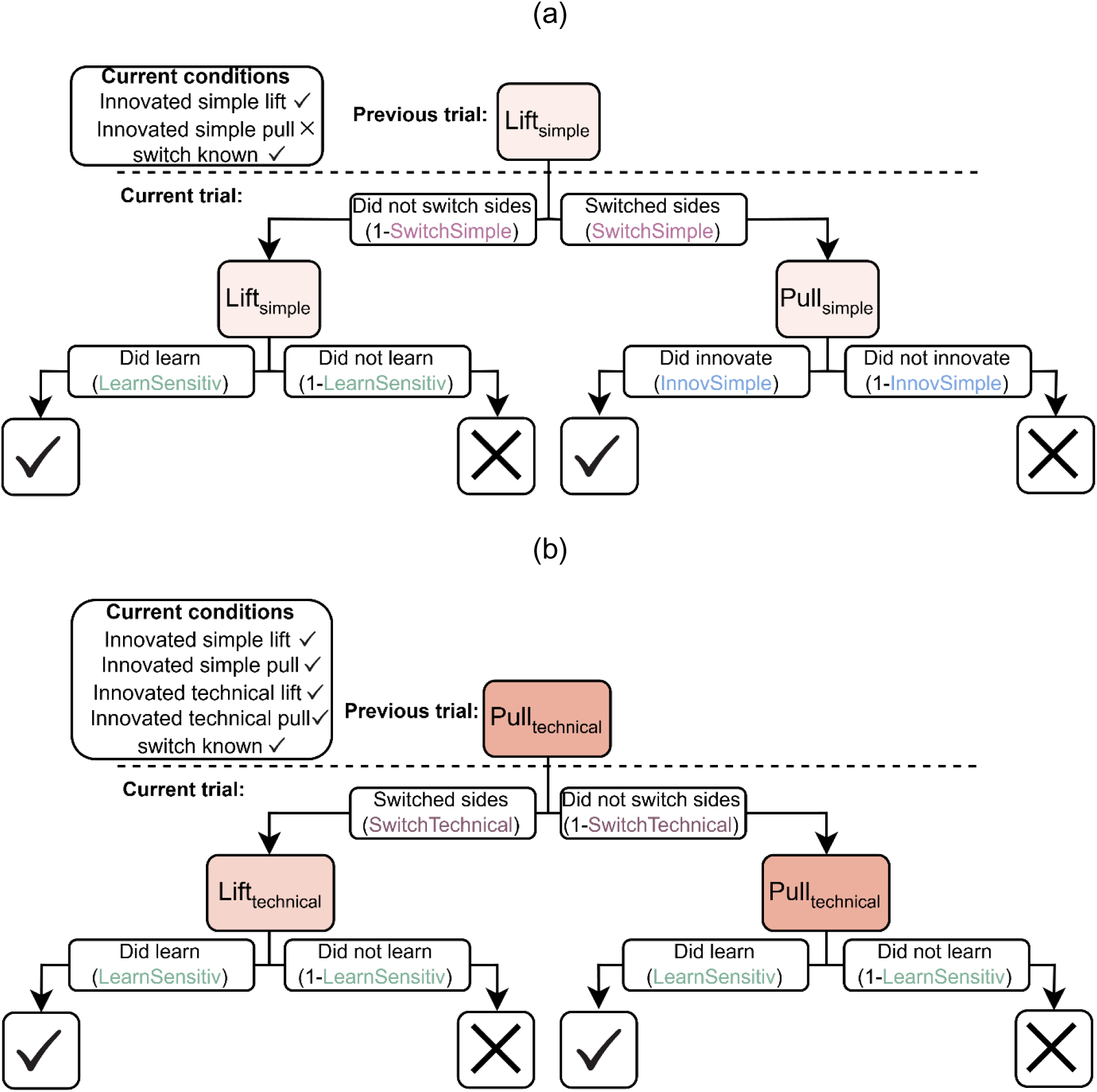
Two examples of decision trees applied to consider the whole temporally ordinated series of each monkey’s interactions with the foraging experiment. The top, opaque box indicates the previous trial choice, the second set of opaque boxes indicates which solution option is attempted, and the bottom set of boxes indicates whether the individual successfully opened the box during this trial (a check means it is successful, a cross means it is a failed attempt). Each branch indicates which parameters are updated if this pathway is taken during the trial. The dashed lines show a separation of trials. (a) The corresponding tree for a phase 1 trial, knowing that the previous trial solution was a lift, so whether there is a switch in this trial is known (“switch” known), the lift solution is known, and the pull solution is unknown. (b) The decision tree corresponding to phase 3 where the previous trial solution was a technical pull solution (“switch” and all solutions are known).

**Table 1.** Estimated parameters for each individual based on their sequence of trial outcomes. The simple parameters represent the performance in phase 1, where the challenge is simpler, while the technical solutions represent the traits expressed when the option of at least one technical solution is present.

| Parameter | Trait | Definition | Phases |
| --- | --- | --- | --- |
| *SS* | Simple Switch<br>Tendency | Switch from the previous trial's solution option to alternative option when both are simple. (1-*SS*) if the individual attempts the same solution again. | 1 |
| *ST* | Technical<br>Switch<br>Tendency | Switch from the previous trial's solution option to alternative option when at least one option is technical. (1-*ST*) if the individual attempts the same solution again. | 2,3 |
| *IS* | Simple<br>Innovativeness | Successfully open the lift or pull for the first time. (1-*IS*) if the individual unsuccessfully attempts the lift or pull solution and has never successfully opened that solution before. | 1 |
| *IT* | Technical<br>Innovativeness | Successfully open the technical lift or technical pull for the first time. (1-*IT*) if the individual unsuccessfully attempts the technical lift or technical pull solution and has never successfully opened that solution before. | 2,3 |
| *V* | Learning<br>Sensitivity | Successfully opened a solution that the individual has previously opened. (1-*V*) if the individual unsuccessfully attempts a known solution. | 1,2,3 |

We assumed that the probability *p* to observe a certain outcome could depend on five parameters constrained between 0 and 1 according to the individuals’ past actions and current phases (table 1): 1. a switch tendency between known simple solutions (“Switch Simple”, *SS*; in phase 1), 2. switch tendency when technical solutions are available (“Switch Technical”, *ST*; in phase 2 and 3), 3. an ability to innovate for the first time one of the simple solution (“Innovate Simple”, *IS*; in phase 1 and 2), 4. an ability to innovate for the first time one of the technical solutions (“Innovate Technical”, *IT*; in phase 2 and 3) and 5. an ability to reproduce previous performance (i.e., a learning sensitivity, *V*). We considered that a monkey switched solutions if they engaged into a solution new for the trial but previously experienced.

We assumed that the monkeys should perform sequential reasoning (44), and thus assumed that *p* was the product of two components: choosing which solution to attempt (lifting or pulling) and the eventual output of the attempt (innovativeness, if the action was never performed before in previous trials, or learning sensitivity, if the action had already been performed in previous trials). For example, if we assumed that an individual *i* in phase 1 at trial *t* has successfully obtained a peanut by lifting, knowing that it had already lifted in an anterior trial but had pulled at trial *t-1*, we would have *p_i,t_* = *SS x V*. This series of decisions, and probabilities of observing a given behaviour, can thus easily be represented by a decision tree (the 45 of them being represented in the electronic supplementary material, figure S2; figure 2 illustrates two examples).

For each monkey, we could estimate the likelihood of the observed trial sequence as the product of all the trial outcome probabilities. We estimated which combination of *SS*, *ST*, *IS*, *IT* and *V* maximized the log-likelihood separately for each monkey using the “optim” function of the *stats* package using the “L-BFGS-B” optimization method to constrain parameter ranges between 0 and 1 (45). Best-fit parameters (and their associated 95% confidence intervals) were used to characterize individuals’ switch tendency, innovativeness and learning sensitivity for further analyses and robustness of our inferences (electronic supplementary material, sections S7 & S8).

#### 2.5.2 Regression modelling and correlations

##### 2.5.2.1 (i): Predicting switch tendency and technical innovativeness

To identify the factors predicting individual variation in switch tendency and technical innovativeness, we modelled individuals’ simple switch tendency scores, *SS*, and technical innovativeness scores, *IT*, as a function of anthropogenic raiding experience, demographic (a composite measure relative to age, sex and rank, electronic supplementary material, section S9) and social composite variables (a composite measure relative to social exposure to the foraging experiment, electronic supplementary material, section S9), to reduce model complexity given the low sample size. To evaluate the respective influence of raiding tendency, social and demographic variables on an individuals’ switch tendency, we used a continuous variable, but for technical innovativeness, we used a binomial regression with a logit error structure (“glmmTMB” function with the family set to “binomial” of the *glmmTMB* package (46)). *IT* was transformed into a binary variable with “low innovativeness” occurring when the confidence interval around the mean *IT* estimate did not cross with 0.5, and “high innovativeness” otherwise (electronic supplementary material, table S7, see figure S13 for a sensitivity analysis of this cut-off choice). All variables were scaled (to a mean of 0 and a standard deviation of 1). We used an information-theoretic approach (47,48) to investigate which variable was likely influential. Specifically, we used the sum of the models’ weight in which the variable of interest was included as a proxy to describe the likelihood that this variable influences innovativeness (with respect to other variables considered) and quantified its average estimate (and associated unconditional confidence interval) by computing the shrunk average model (i.e. considering all models and their weights; “modavgShrink” function of the *AICcmodavg* package (49)). We ensured that the most complex model fit was good by (1.) verifying that all models converged and were not singular (“check_convergence” and “check_singularity” functions of the *performance* package (50)), (2.) visually and statistically checking the models’ assumptions (i.e. adequate distribution of residuals under the modelled distribution and their homoscedasticity using the *DHARMa* package (51); electronic supplementary material, figures S14 and S15), (3.) verifying the absence of major outlier (“check_outliers” function of the *performance* package (50)), and (4.) verifying the absence of over-parameterisation (“extractCN” function of the *AICcmodavg* package (49); all parameters had low condition values, electronic supplementary material, figure S14). We also verified that there was no collinearity issue by focusing on the Variance Inflation Factor (VIF < 2, “check_collinearity” function of the *performance* package (50), electronic supplementary material, tables S8 and S9).

##### 2.5.2.2 (ii): “Behavioural flexibility syndrome”

To investigate the link between multiple traits potentially underlying anthropogenic foraging in vervet monkeys, we evaluated whether the three traits *IT, SS,* and *V* formed a behavioural flexibility syndrome (i.e., a group of correlated behavioural traits (52)). To do so, we modelled each variable as a function of the other using beta-regression (as the variables are constrained between 0 and 1 (53)) with a logit error structure (“glmmTMB” function with the family set to “beta_family” of the *glmmTMB* package (46)). Prior to fitting, we transformed each output variable *y* such as *y’ = (y(n − 1) + 0.5)/n*, where *n* represents the sample size, in order to account for values strictly equal to 0 or 1 (54). As our focus was on individual problem-solving skills that required flexibility, as a measure of innovativeness and switch tendency, we modelled *SS* and *V* as a function of *IT*, and *V* as a function of *SS*.

##### 2.5.2.3 (iii) Predicting problem-solving and human food consumption

We first evaluated whether experimental performance links to a monkey’s human food consumption in the urban habitat using a Pearson correlation (“cor.test” of the *stats* package (55)).

We then separately investigated the drivers of two measures of success (in the foraging experiment and in exploiting human food). For monkeys to successfully retrieve the food reward in phase 3 of the experiment, they were required to use their motoric and technical skills, as well as to innovate the solution to the problem presented to them, either innovating the technical lift or technical pull. To avoid biasing the technical innovativeness scores, we excluded phase 3 from our foraging experiment success calculations. We tested the effects of individuals’ switch tendency (approximated by *SS*) and innovativeness (approximated by *IT*) on predicting foraging success in phase 1 and 2. We considered the foraging experiment success rate for each individual *i* across the two phases, *s_i_* as the weighted average of success rate between the two phases such as *s_i_ = (1-w)s_i,1_ + ws_i,2_* with *w* = *s_1_/(s_1_*+*s_2_*) where *s_1_* and *s_2_*represent the success rate of all the monkeys tested in phase 1 and 2 respectively. We then modelled the foraging experiment success rate (constrained between 0 and 1) as a function of *IT* and *SS* with a beta-regression (implemented as in 2.4.2.1), considering *SS* with a second-order polynomial form. We used a second-order polynomial (*SS^2^*) based upon the reasoning that a monkey could be persistent (low *SS* and thus using same solution over and over) or highly flexible (high *SS*) to successfully retrieve the reward.

To allow for estimate comparability (56), both continuous predictors were scaled to a mean of 0 and a standard deviation of 1. We then repeated this procedure with human food consumption as the response variable. We verified the model fit quality as in 2.5.2.1 (electronic supplementary material, tables S10 and S11 and figures S16 and S17).

## 3. Results

Out of the 42 monkeys from both troops, 41 were initially tested (only one did not engage in the experiment) and 80.5% completed at least 10 trials in phase 1 (33/41, N_phase1_ = 444). Of these, 81.8% made it to phase 2 (27/33, N_phase2_ = 174) and 92.6% of those individuals made it to phase 3 (25/27, N_phase3_ = 155). The number of trials per monkey per phases 1, 2, and 3 were 12.8 (± 3.64), 6.40 (± 1.03), and 6.5 (± 1.27), respectively (when not indicated otherwise, we always refer to a variable by its mean ± sd.). For the three phases, the overall success rate was 87.2% (387/444), 81.6% (142/174) and 41.3% (64/155), respectively.

27 monkeys (65.9%) performed sufficient technical trials to characterize their problem-solving skills in terms of switch tendency between simple (*SS*) and technical solutions (*ST*), innovativeness (either simple *IS* or technical *IT*), and learning sensitivity (*V*). Individuals’ estimates are presented in figure S12. Overall, switch tendency was relatively low (*SS*: 0.263 ± 0.032; *ST*: 0.469 ± 0.036) and innovativeness varied across technical levels (69.6% and 11.1% with *IS* > 0.5 and *IT* > 0.5, respectively). When the monkeys successfully performed a solution, they were generally good at reproducing it (*V*: 0.792 ± 0.025).

### 3.1 Anthropogenic experience did not predict innovativeness in urban monkeys

The tendency to switch between the lift and the pull options (phase 1) was not explained by demographic traits, previous social exposure, or a monkey’s raiding tendency (*null* model weight: 0.64, electronic supplementary material, table S5). The ability of monkeys to innovate the technical solution was best explained by demographic and social exposure rather than the tendency to raid anthropogenic structures (figure 3, left). While the effect was largely uncertain (hence not significant, as expected given the reduced sample size), the demographic trait component showed the strongest effect, est. [CI_95%_] = 0.388 [-0.463, 1.239], followed by the social component, est. [CI_95%_] = 0.118 [-0.479, 0.715], and raiding tendency rates explained the least of the technical innovativeness scores, est. [CI_95%_] = -0.003 [-0.386, 0.392] (figure 3, right). Note that the *null* model (not accounting for any of these factors), was nonetheless not part of the 95% confidence set of the best models (table 2), suggesting that these factors were likely influential altogether.

**Figure 3.**
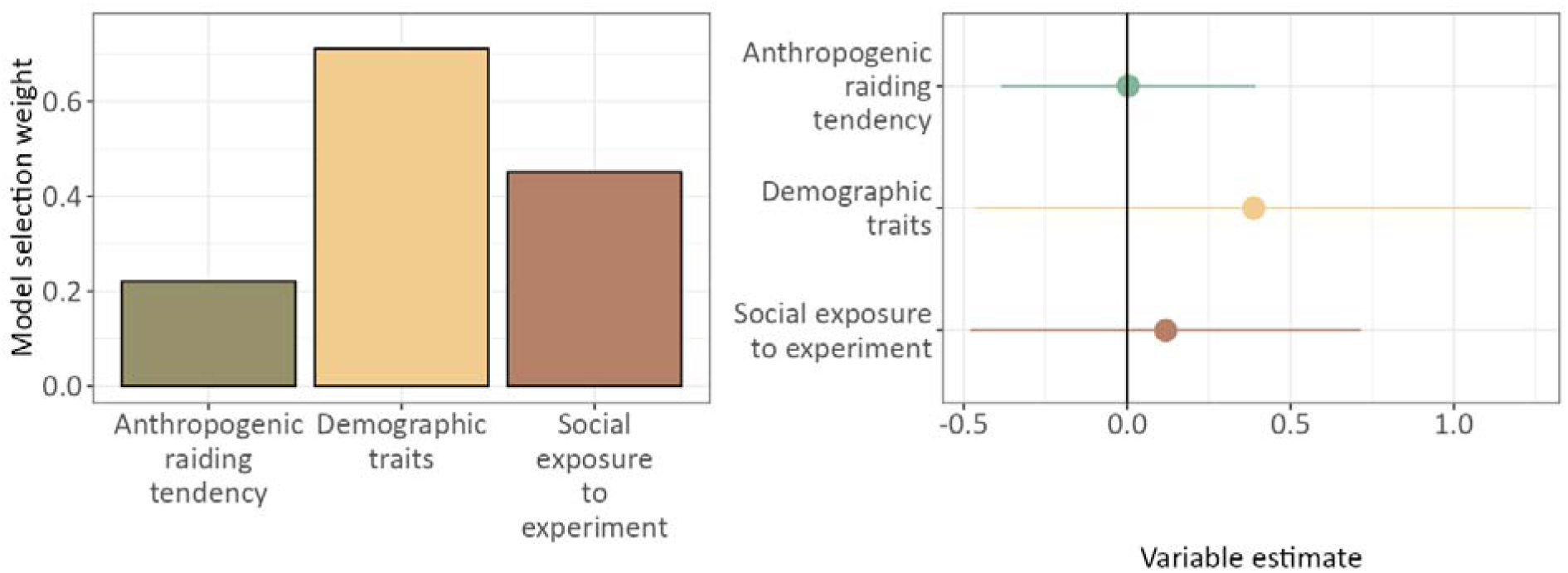
The probability that a monkey solved the foraging experiment through a technical innovation was more likely explained by demographic traits and previous social exposure to the experiment, rather than anthropogenic raiding tendency. (Left) Weights are calculated based on the sum of the AIC weights for every model that includes the respective dimension. (Right) The points indicate the effect estimate and the lines indicate the associated 95% confidence interval.

**Table 2.**
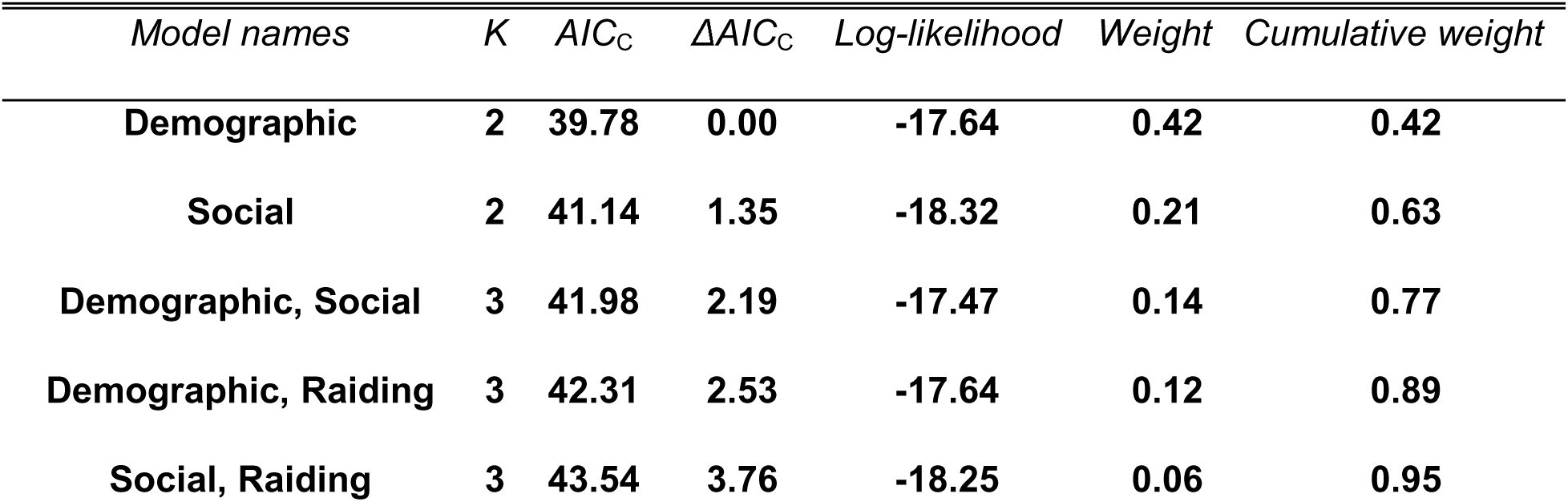
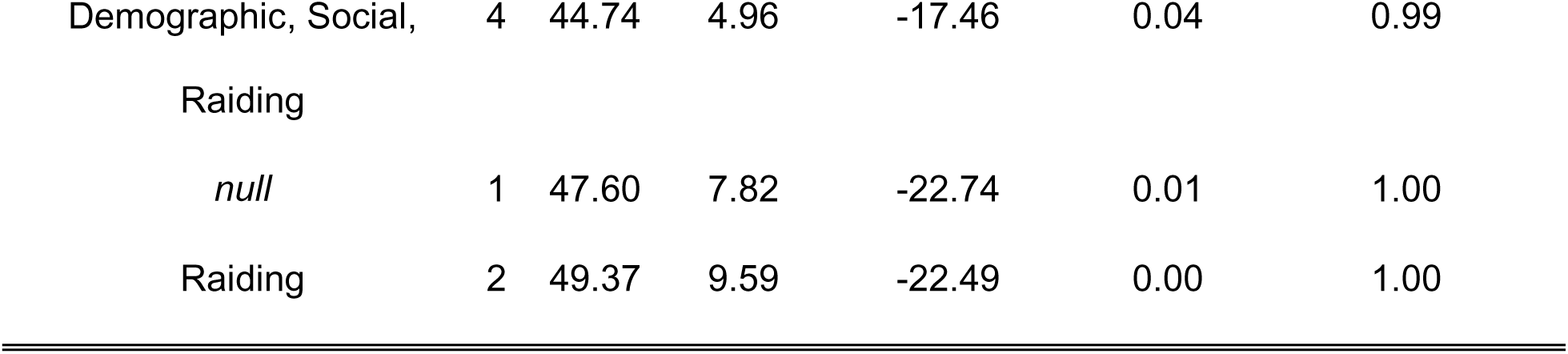
Summary of the model selection approach. *K* represents the number of estimated parameters for each model, *AIC_C_*is the Akaike Information Criterion corrected for small samples (Burnham et al. 2011), Δ*AIC_C_* is the difference with the “best” model (demographic only model). Bold rows highlight models constituting the 95% confidence set.

### 3.2 Absence of a behavioural flexibility syndrome

Monkeys that were the best innovators were also the ones most capable of performing a solution subsequently (beta-regression linking *IT* and *V*: est. [CI_95%_] = 1.688 [0.226, 3.150]; electronic supplementary material, table S12). Those innovators were not, however, the monkeys that switched between solutions the most (beta-regression linking *IT* and *SS*: est. [CI_95%_] = -0.863 [-2.577, 0.850]). In addition, switch tendency was unrelated to learning sensitivity (beta-regression linking *SS* and *V*: est. [CI_95%_] = -0.727 [-2.104, 0.651]).

### 3.3 Innovativeness predicted problem-solving skills but not human food consumption

Foraging experiment success did not correlate with human food consumption (ρ_pearson_ [CI_95%_] = 0.099 [-0.253, 0.428], t_31_ = 0.553, p = 0.584). *IT* significantly predicted monkeys’ performance in the foraging experiment (est. [CI_95%_] = 2.14 [0.447, 3.83]; table 3) but not *SS* (X_1_^2^ = 1.036, p = 0.596). Neither *IT, SS, nor SS^2^* predicted an individual’s human food consumption (*null* vs. *full* model comparison: Likelihood Ratio Test: X_3_^2^ = 1.828, p = 0.609; table 3 and figure 4).

**Figure 4.**
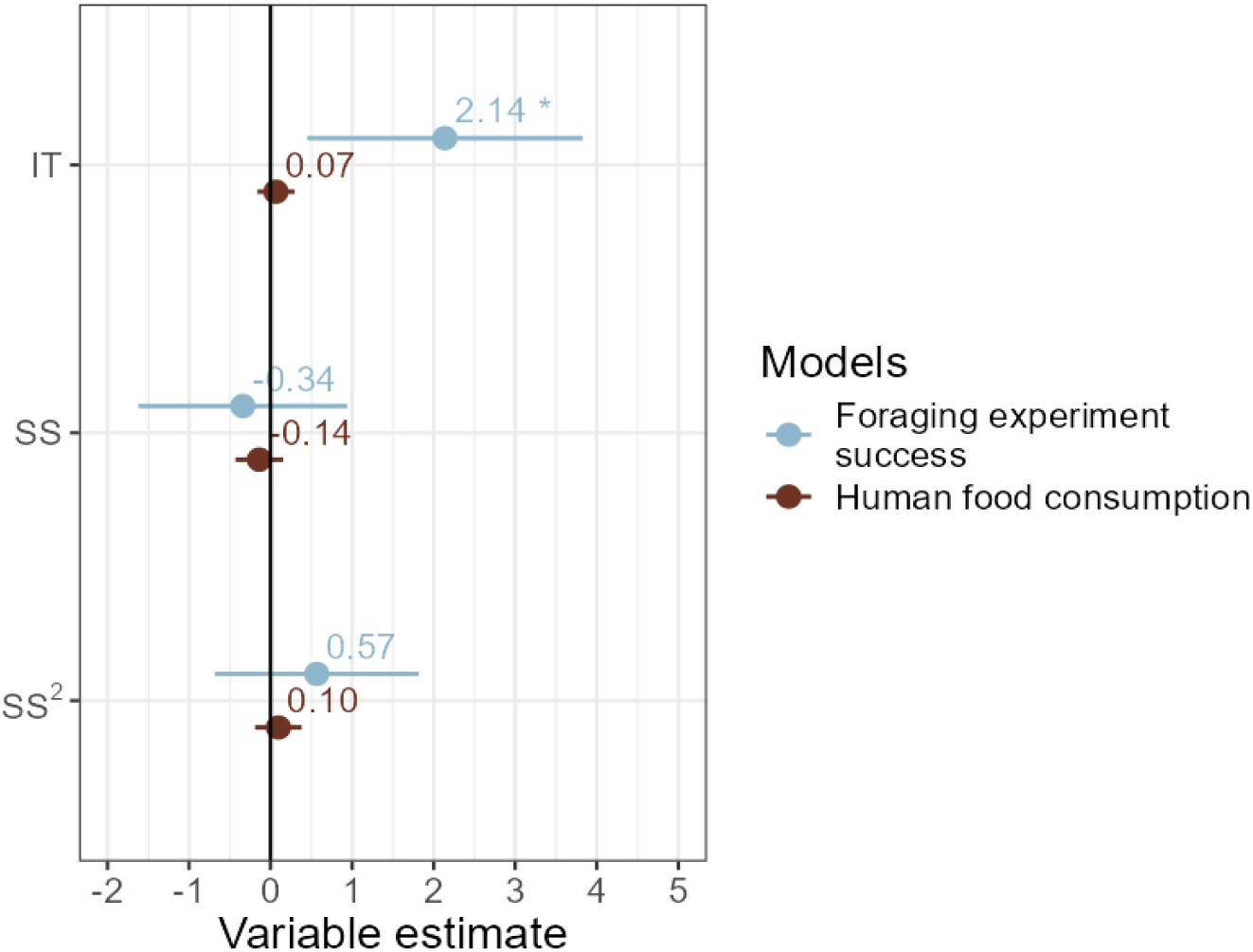
Forest plot of the beta-regression of individuals’ success in the foraging experiment and human food consumption as a function of the switch tendency between simple solutions (polynomial term, *SS^2^* + *SS*) or the capacity of innovating a technical solution (*IT*). The points indicate the estimate, and the lines indicate the associated 95% confidence interval. The estimate values are given above each point and the “*” indicates the p-value significance.

**Table 3.**
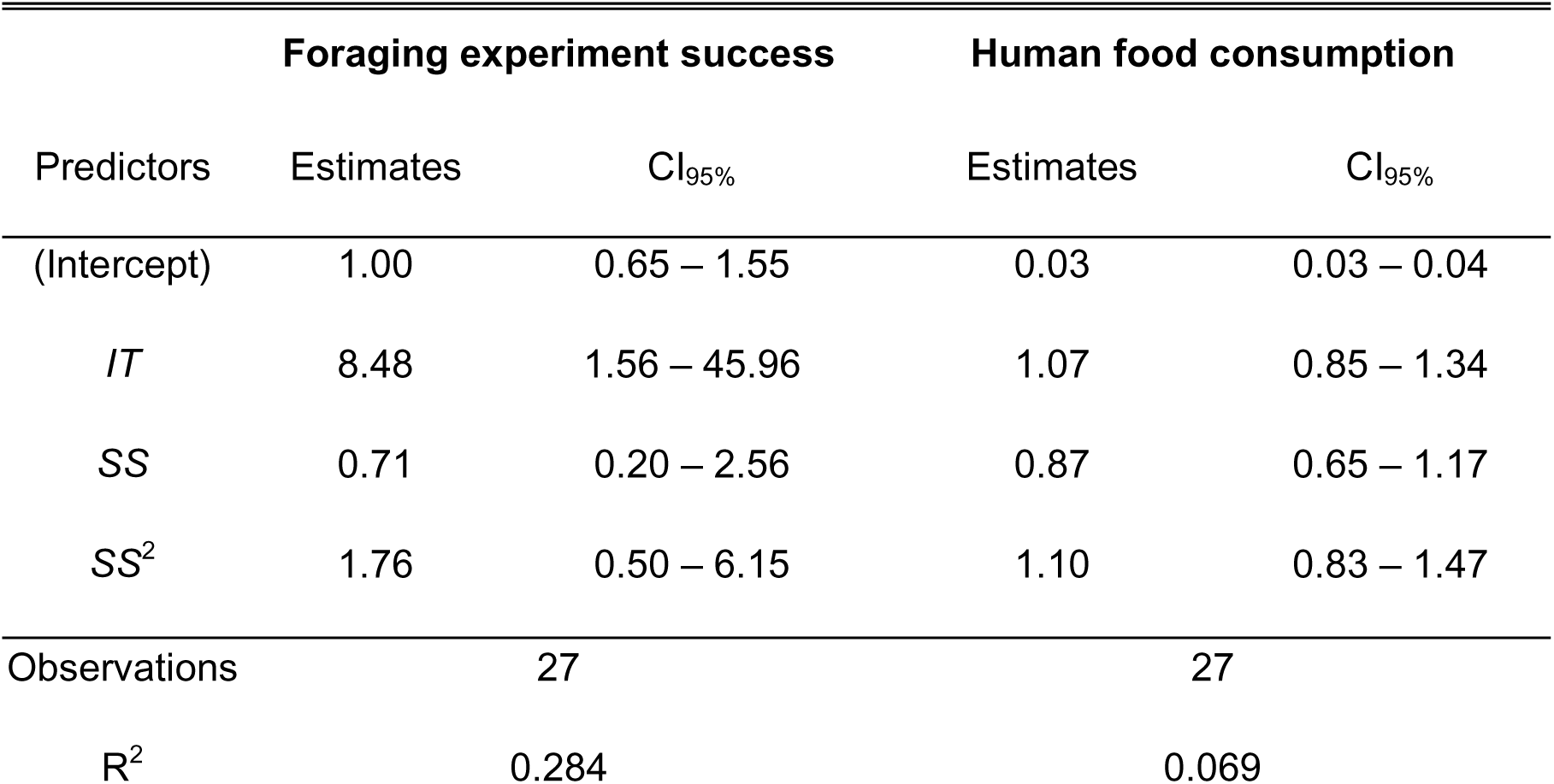
Summary of success measure models. CI_95%_ = Confidence interval at the 95% level. The estimates are raw estimates (i.e., not adjusted for the logit transformation).

## 4. Discussion

Our study investigated the link between anthropogenic raiding tendency and behavioural flexibility (switch tendency, innovativeness, and learning sensitivity) using a foraging experiment with semi-urban vervet monkeys. Participation in the task was high: over 80% of initial participants completed phase 1, providing a robust and representative sample of the population. We found, first, that monkeys with higher rates of anthropogenic foraging experience did not appear more behaviourally flexible, neither in their switch tendency, nor in their innovativeness. Second, the expected behavioural flexibility syndrome composed of these traits was not supported by our data. Third, behavioural flexibility did not predict monkeys’ human food consumption.

### Exaptation rather than adaptation

None of the three factors (raiding tendency, social exposure or demographic traits) predicted switch tendency, and raiding tendency was less predictive of technical innovativeness than demographic traits or previous social exposure to the puzzle box (table 2, figure 3). The influence of social exposure aligns with previous research in vervet monkeys, which underscores the importance of social learning in the acquisition and spread of problem-solving behaviours (32,33,57).

As raiding tendency was not a significant predictor, these results suggest that the traits observed in our foraging experiment may fall within the “latent solution” of cognitive capacities for vervet monkeys as a species (22) rather than representing skills shaped by their urban experience. That is, the behavioural flexibility underpinning problem-solving may be species-wide, pre-existing traits evolved for different challenges, such as extractive foraging, rather than adaptations to urban life. Indeed, vervet monkeys being dietary generalists use their behavioural flexibility while foraging in natural environments where they switch between food items, e.g. from eating leaves up in trees to opening hard seedpods, licking nectar from flowers or rolling stones on the ground to access hidden insects.

Urban environments are often assumed to select for enhanced behavioural flexibility due to the presumed novelty of urban challenges (58). Since raiding frequently involves the manipulation of human-made objects to access food, we expected that individuals who raid more would show higher levels of behavioural flexibility when interacting with our novel foraging task, but this was not the case. Our study population of vervet monkeys have experienced this semi-urban environment throughout multiple generations and certainly the entire lives of the participating monkeys. For these individuals, anthropogenic elements, such as man-made structures and objects like bins with lids do not represent novel challenges. This familiarity may alleviate the selection pressure for enhanced innovativeness. Instead, urban living may permit exaptation, where the expression of ancestral traits evolved for one function are co-opted for another (59), allowing urban monkeys to respond to familiar, albeit anthropomorphized, environments. Our empirical findings align with theoretical suggestions that urbanization does not necessarily require increased innovativeness, but rather a flexible application of pre-existing cognitive capacities such as the ability to learn about the urban environment (60). Aligning findings by Vardi and Berger-Tal (61) show that behavioural flexibility of male house sparrows (*Passer domesticus*) is better predicted by the rate of anthropogenic change rather than its magnitude, suggesting that, in relatively stable urban environments, animals may not require novel cognitive adaptations to persist.

Experimental studies of cognitive traits in urban primates remains limited (42,62–64). Of these, only Pal et al. (62) directly linked a metric of anthropogenic exposure to task performance in bonnet macaques (*Macaca radiata)* and found a positive correlation between locals and tourists actively feeding the monkeys and the frequency of monkey’s sophisticated bottle opening techniques, although food provisioning may have confounded experience with motivation. In contrast, our data showed no such correlation, as raiding did not predict innovativeness nor switch tendency. As these differences occur, this highlights the possibility that these motivational differences also affect the interpretation of rural and urban population comparison studies. By examining individuals within the same population, we controlled for environmental differences in motivation, highlighting the value of within-population approaches when evaluating the effects of urban experience on cognition.

### Flexibility traits did not form a behavioural syndrome

Using a probabilistic modelling approach, we included the entire sequence of all three experimental phases and trials to trace an individual’s behavioural actions and determine their individual switch tendency, innovativeness, and learning sensitivity. This provided a more robust reflection of the monkeys’ cognitive reasoning compared to trial averages or endpoint scores alone. Although previous work applying similar methods used binary outcomes and more trials (65), our model validation (electronic supplementary material, section S8) showed robust outcomes and a lack of sensitivity to arbitrary decisions we had to make.

Evolutionary theory suggests that behavioural syndromes, or correlated sets of traits, can evolve when feedback loops reinforce co-expression (66). For example, in some species, boldness and aggression correlate because bolder individuals secure more resources, which further enhances aggressiveness (66). If urban environments imposed such consistent selection on traits underpinning behavioural flexibility, we might expect stronger trait integration. Contrary to our predictions, we found no relationship between the three flexibility traits (switch tendency, innovativeness and learning sensitivity), except for a positive relationship between technical innovativeness and learning sensitivity (section 3.2). Switch tendency was not associated with either of these. Both technical innovativeness and learning sensitivity are processes likely underpinning solving of a more challenging physical problems (as posed in phase 2 and 3), whilst switch tendency possibly relates more to motor actions used by monkeys frequently, especially in the urban habitat. This suggests that switch tendency may reflect an exploratory tendency or motivational trait, rather than an advanced cognitive capacity. These findings are consistent with prior work showing that behavioural flexibility measured by switching between options was not correlated with learning in grey squirrels (1).

The absence of an adaptive, behaviourally flexible syndrome suggests that selection may not uniformly act upon switch tendency, innovativeness, and learning sensitivity. Just as modelled mixed-ability groups (e.g., diverse problem-solving agents) can outperform homogenous groups of the high-ability problem solvers (67), variation in motivational traits and flexibility among troop members may buffer populations against a range of urban challenges. In vervet monkeys, traits such as boldness or risk-taking may interact with, or even overshadow, cognitive flexibility in determining urban success.

### Flexibility traits relating to problem-solving is not related to human food consumption

Innovativeness, but not switch tendency, significantly predicted individual success in the foraging experiment (table 3, figure 4), which aligns with comparative findings across bird species (68). Conversely, in our study, neither trait predicted an individual’s consumption of anthropogenic food sources. This was unexpected, given that human food consumption represents attractive, opportunistic, high calorie foraging opportunities that benefit individuals (at least in the short term), and during which monkeys encounter human artefacts and as such occasionally may need to innovatively solve technical problems. Though, at our study site of Simbithi Eco-Estate, vervet monkeys still forage primarily on natural vegetation (34) and therefore, the vast abundance of forest patches likely buffers any potential effects of urbanization for their need to exploit human food sources, and thus necessity for technical innovativeness (69).

Since it is difficult to obtain classic fitness measures to quantify success in this environment, such as longevity and reproductive success directly in long-lived species like primates (70,71), we used human food consumption as an approximation of success in habitat exploitation. However, this proxy did not relate to any of our behavioural flexibility measures. Foraging success, especially risky ones involving entering human houses, may be mediated by personality traits or social factors. For instance, dominant individuals may monopolize human retrieved food or benefit through sharing from other bold individuals’ efforts to take such risks (72,73). Additionally, vervet monkeys are known to socially learn their dietary preferences (57,74), which may further reduce selection on individual flexibility and contribute to the observed missing link between innovation and human food consumption in our data. Consequently, social relationships may be more decisive for food tolerance and the development of food preferences and foraging skills. To sum up, our results suggest that behavioural flexibility, while beneficial in a controlled foraging experiment, is likely not decisive of success in semi-urban environments.

### Limitations and broader implications

Although our results suggest that current levels of behavioural flexibility in vervet monkeys suffice for them to thrive in the current anthropogenic environment, our measurements represent only a snapshot in time, and they do not rule out future adaptative change. While most features specific to the anthropogenic habitat are independent from the monkeys’ presence, the human inhabitants are often using targeted prevention measures to hinder monkey raiding (i.e. monkey screens to prevent access to houses, bin clips, spinning bin tops). As such, monkeys may be involved in an arms race with the humans (like between cockatoos (*Cacatua galerita*) and humans, (30), applying selective pressures affecting monkeys’ cognitive profiles. As Simbithi Eco-Estate was established in 2003 (22 years ago), the selection pressures may change as prevention methods increase and the eco-estate matures, potentially leading to increased human-wildlife conflicts (75). Re-testing this population of monkeys after several years may reveal changes in the population’s behavioural flexibility. Combining this with the known life-histories and pedigrees of the monkeys in the future will lead to a better understanding of how the urban environment shapes animal cognition, and may help develop management strategies to limit human-vervet conflicts.

## Author contributions

<u>Paige Barnes</u>: conceptualization, data curation, formal analysis, investigation, software, methodology, validation, visualization, writing – original draft preparation, writing – review & editing

<u>Benjamin Robira</u>: conceptualization, supervision, data curation, formal analysis, methodology, software, validation, writing – original draft preparation, writing – review & editing

<u>Stephanie Mercier</u>: investigation, supervision, project administration, data curation, writing – review & editing

<u>Sofia Forss</u>: conceptualization, funding acquisition, resources, supervision, project administration, writing – review & editing

## Acknowledgements

We would like to sincerely thank the Simbithi Environmental board for their support to the Urban Vervet project. We also thank the KONE Foundation (Finland; SF), SNSF Grant number CRSK-3_220769 (SF) and the University of Zurich’s Fonds zur Förderung des Akademischen Nachwuchses and the Gebauer Stiftung (Switzerland; SF) for their financial support of our work. Thank you to Melissa Ardila Villamizar, Natacha Bande, Emma Chen, Manon Desaivres, Lindsey Ellington, Joey Felsch, Oceane Lusher, Zonke Mbutho, Adrian McConnell, and Ntaki Senoge for their contributions to the behavioural observation dataset. Furthermore, we thank the iNkawu Vervet Project & Charlotte Canteloup for lending their puzzle boxes to us and Tim Driman for his help with modifications. Thank you to Lily Johnson-Ulrich for manuscript advice.

## Data and code availability

Data can be available for review upon request to the corresponding author. The codes used to analyse these data are available for review at https://github.com/paigebarnes/BehaviouralFlexibilityVervet. Both data and codes will be made available in a permanent repository upon acceptance.

