## Supplementary Material for "Behavioural flexibility and its drivers in semi-urban vervet monkeys"

*1. Trial visualizations*

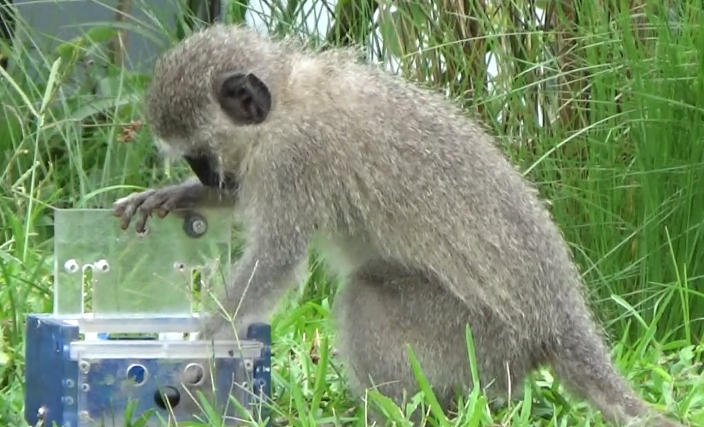

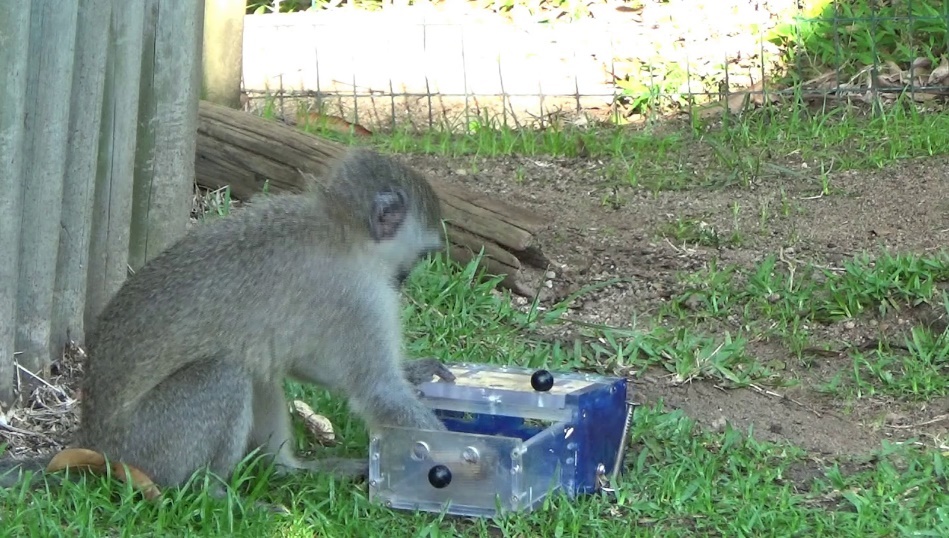

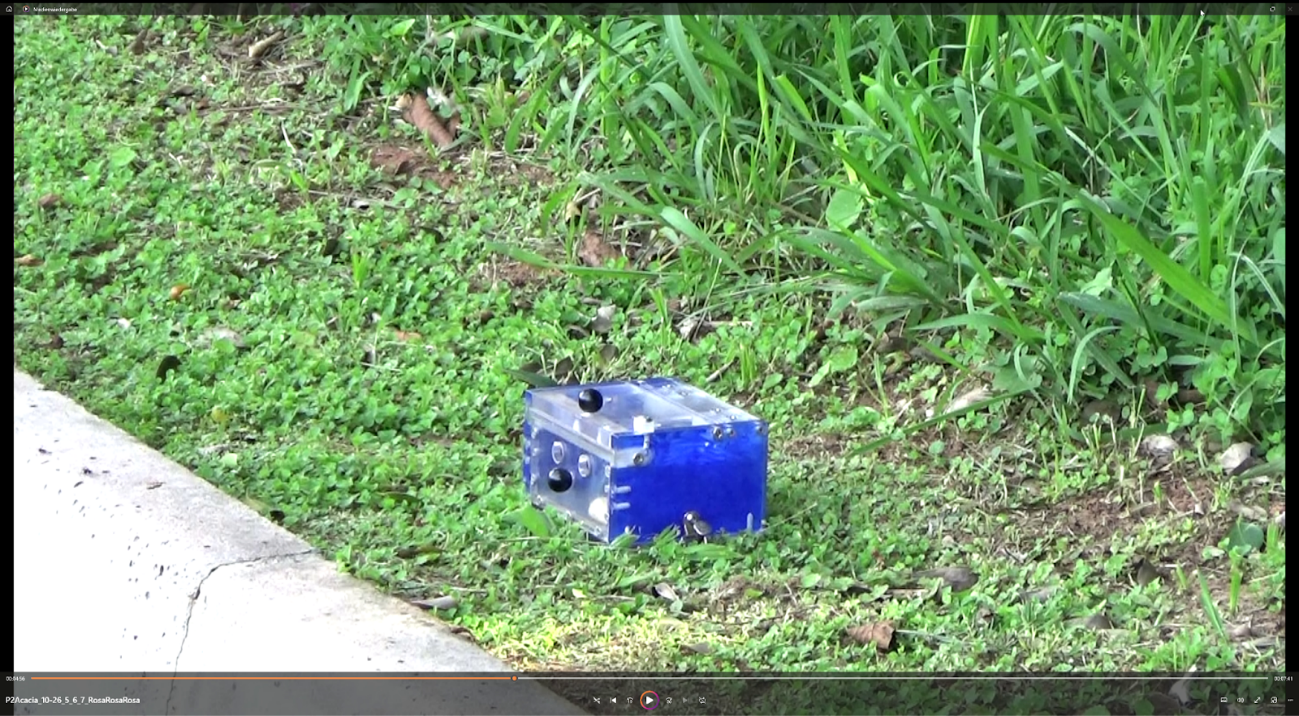

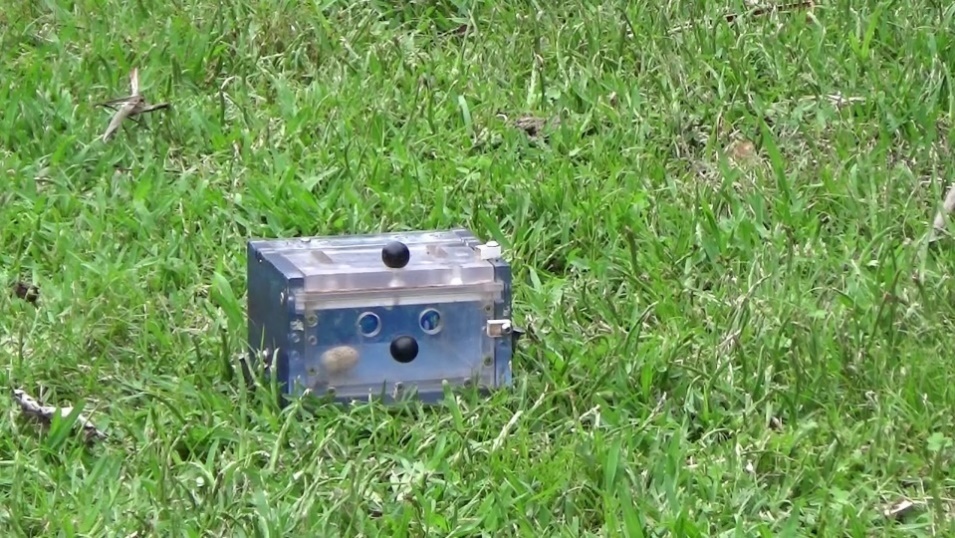

**d)**

**c)**

**b)**

**a)**

**Figure S1.** Example photos for each possible box solution across all three phases. The phase 1 simple solutions are a) lift and b) pull. Phase 2 included the b) pull option, but also c) a lock on top for the lift solution, making this a technical lift. Phase 3 had the c) technical lift and the d) technical pull, which included a lock on the side as well.

Video examples for each phase are available online:

 
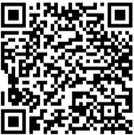

*2. Experimental ethogram*

**Table S1.** Ethogram used when coding videos of the experiments to determine the outcome of each puzzle-box trial.

| **Outcome** | **Behaviour** |
| --- | --- |
| Lift success | The monkey manipulates the box so that the peanut is accessible by the subject through the lift solution. This is usually done by lifting the round knob on the front of the lid, or the lid itself, upwards (figure 1a). |
| Pull success | The monkey manipulates the box so that the peanut is accessible by the subject through the pull solution. This is usually done by pulling open the round knob or pushing open from the back of the box (figure 1b). |
| Lift attempt (unsuccessful) | The monkey touches or bites the top of the box and/or lift the knob, with some movement (grabbing, pushing). The subject should be looking at the box while interacting with this side. |
| Pull attempt (unsuccessful) | The monkey touches or bites the front or back of the box and/or pull knob, with some movement (grabbing, pushing). The subject should be looking at the box while interacting with these sides. The back is included because the pull can be pushed open from the back. |

*3. Trial outcomes for individual monkeys*

**Table S2.** Trial outcome for phases 1 to 3.

**Phase 1**

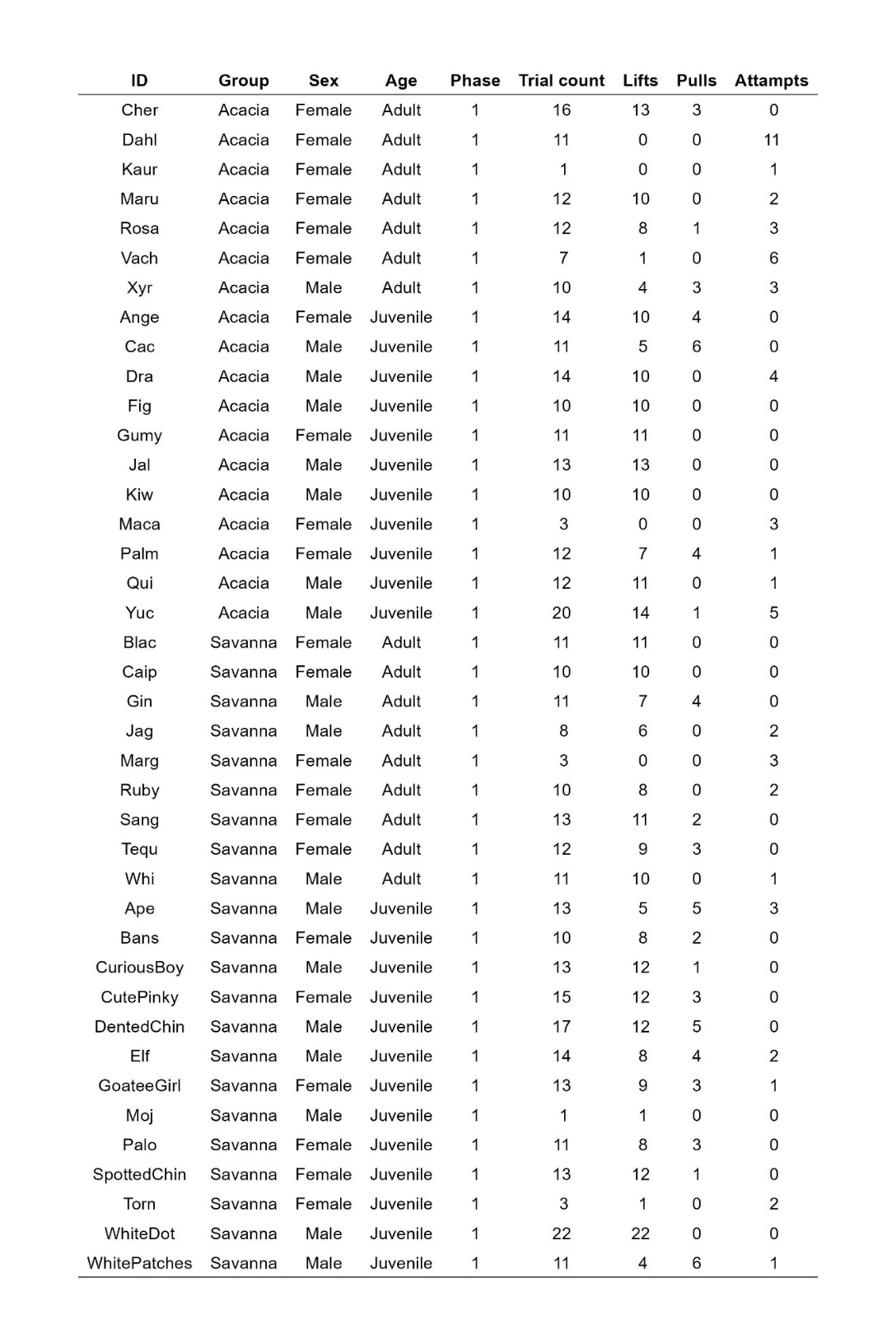

**Phase 2**

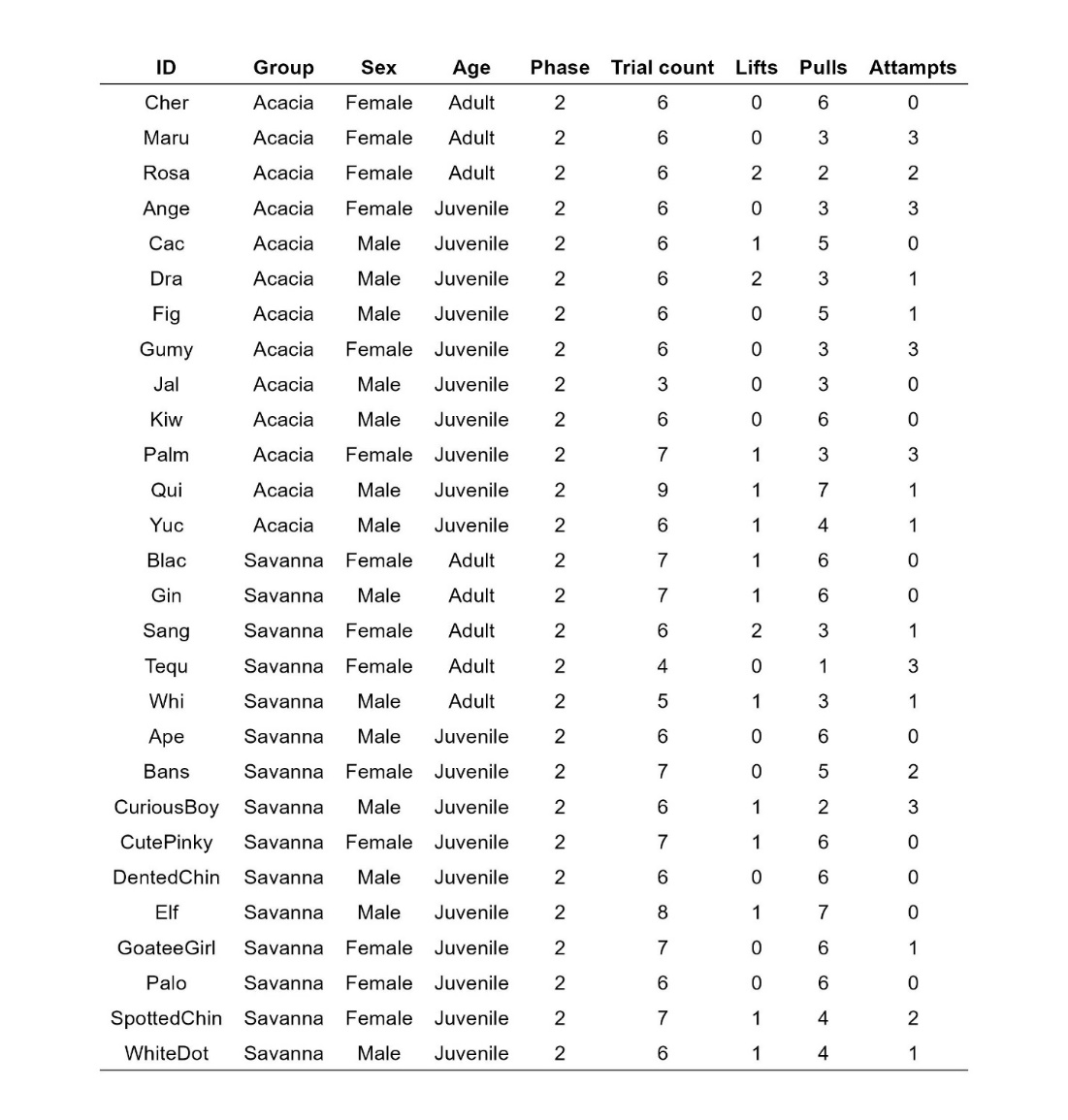

**Phase 3**

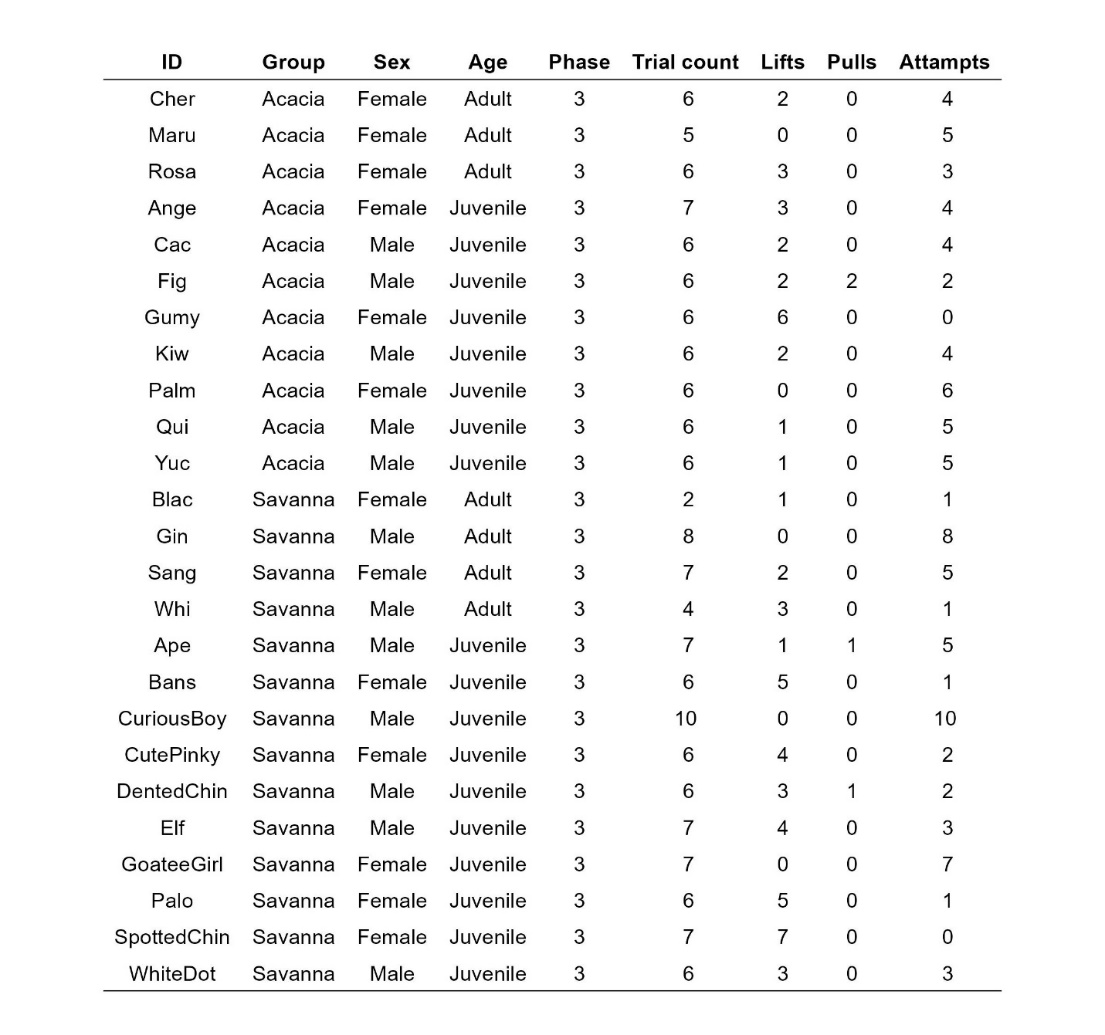

*4. Behavioural observation ethogram*

**Table S3**. Ethogram used when collecting ad-libitum behaviours.

| **Category** | **Behaviour** | **Description** |
| --- | --- | --- |
| Agonistic | Attack | Individual moves forward or lunges to an opponent. |
|  | Stare | Individual lifts out its eyelids (white/pink above its eyes is visible) while looking intensely at another monkey, usually combined with attack. |
|  | Broadside display | An individual stretches, flexes, and stays “unnaturally” stationary in front of another, usually performed by males towards males in a dominance context. |
|  | Bite | Individual bites another one. |
|  | Grab | Individual grabs another one. |
|  | Head-bob | Individual is moving the head up and down repeatedly. |
|  | Hit | Individual hits another one. |
|  | Sit on others | Individual approaches and sits on another one who might vocalize and wait for the other to leave. Probably most often in an agonistic context when a high-ranking individual sits on a lower-ranking one. |
|  | Take place | Individual displaces another and takes its place, usually combined with an approach. |
|  | Chase | Individual runs after another one who is fleeing. |
|  | Fight | When two or more individuals engage in an aggressive interaction, so intense and fast that it is impossible to disentangle the behaviours and the identity of each specific action. |
|  | Vocalize | Individual produces any call that is not an alarm call or a scream. |
|  | Yawn | Individual yawns while showing their teeth. This behaviour only applies in agonistic contexts. |
|  | Stand-up | Individual stands up on feet, usually during an agonistic event. |
|  | Steal food | Individual takes another individual’s food. |
|  | Frustration hop | Individual jumps up and down, usually while screaming and potentially looking intensely at another individual. |
| Submissive | Crawl | Individual bows down to an aggressor while looking at it. |
|  | Flee | Individual runs away from an aggressor who is chasing. |
|  | Jump aside | Individual jumps aside to avoid another one. |
|  | Retreat | Individual quickly leaves the proximity of another aggressive one (displacement but not pursued as in a chase). |
|  | Avoid | Individual moves its head or body away from an aggressor to avoid an interaction (no displacement). |
|  | Teeth-chattering | Individual makes rapid jaw movements that are noisy, usually between males of different ranks, to initiate close proximity/grooming. |
|  | Look around | Individual looks around at targeted individuals in the vicinity. |
|  | Scream | Individual produces a distress call/scream usually during agonistic interactions. Note: screams can be produced by both victims & aggressors. |
| Anthropogenic interaction | Raiding | A monkey interacts with a house, garage, bin, bag, car, braai, or bird feeder. |
|  | Anthropogenic feeding | The monkey ingests food that is not found in the natural habitat (bread, apples, etc) or taken from houses or garbage. |

*5. Individual identification test and inter-observer reliability tests*

To ensure that all data were collected in a standardized way, each observer had to pass both an individual identification test on each monkey of their respective studied troop (ID test), as well as an inter-observer test on both scan and focal data with the project manager. In an ID test, observers had to correctly identify each monkey consecutively within 30 seconds under at least three out of four conditions: *easy*: individuals nicely visible within 10 m; *high*: individuals higher than observers up in a tree, *far*: individuals at a distance greater than 10 m away and *difficult*: individuals being either far and high, or obscured in a dense vegetation patch.

For observers to pass behavioural inter-observer reliability tests, a mean proportion of agreement of at least 80% had to be reached for both scan data and focal follows. For the scans, a minimum of 50 data points collected simultaneously by both the observers and the project manager had to be collected and the agreement between observations was calculated (table S4). The agreement between the observers was estimated using the Cohen’s Kappa for two raters using the “kappa2” function of the *irr* package, with the unweighted option for all relevant categories (Gamer et al, 2012).

For the focal follows, a minimum of at least three individuals had to be followed and data collected continuously for 20 minutes on each focal individual by both the tested observer and the project manager simultaneously (one focal on an adult male, one on an adult female, and one on a juvenile). Proportions of agreements have been calculated using the intraclass correlation coefficients for the following categories: number of total data collected, presence/absence of baby for mothers, main behaviour, more detailed behaviour type, and habitat type (“icc” function of the *irr* package, Gamer et al, 2012).

**Table S4**. Results of inter-observer reliability tests between the project manager (PM) and each observer involved in the dataset. The cutoff for reliability increased from 70% to 80% after the first set of observers for the scan data.

|  | Scan Count | Cohen’s Kappa Agreements | Focal Count | IntraClass Correlation Coefficient Agreements |
| --- | --- | --- | --- | --- |
| PM-Observer 1 | 105 | 70% | 4 | 93% |
| PM-Observer 2 | 74 | 73% | 5 | 97% |
| PM-Observer 3 | 91 | 84% | 3 | 96% |
| PM-Observer 4 | 67 | 80% | 4 | 95% |
| PM-Observer 5 | 75 | 80% | 6 | 98% |
| PM-Observer 6 | 83 | 80% | 5 | 99% |
| PM-Observer 7 | 60 | 86% | 7 | 99% |
| PM-Observer 8 | 50 | 81% | 3 | 96% |
| PM-Observer 9 | 101 | 81% | 4 | 96% |
| PM-Observer 10 | 51 | 83% | 4 | 99% |
| PM-Observer 11 | 133 | 82% | 8 | 89% |

*6. Modelling vervet monkeys’ decision steps during the puzzle-box experiment*

We present in figure S2 here below all the decision trees for each phase which were used to estimate the probability of an individual performing a certain behaviour.

**Phase 1**

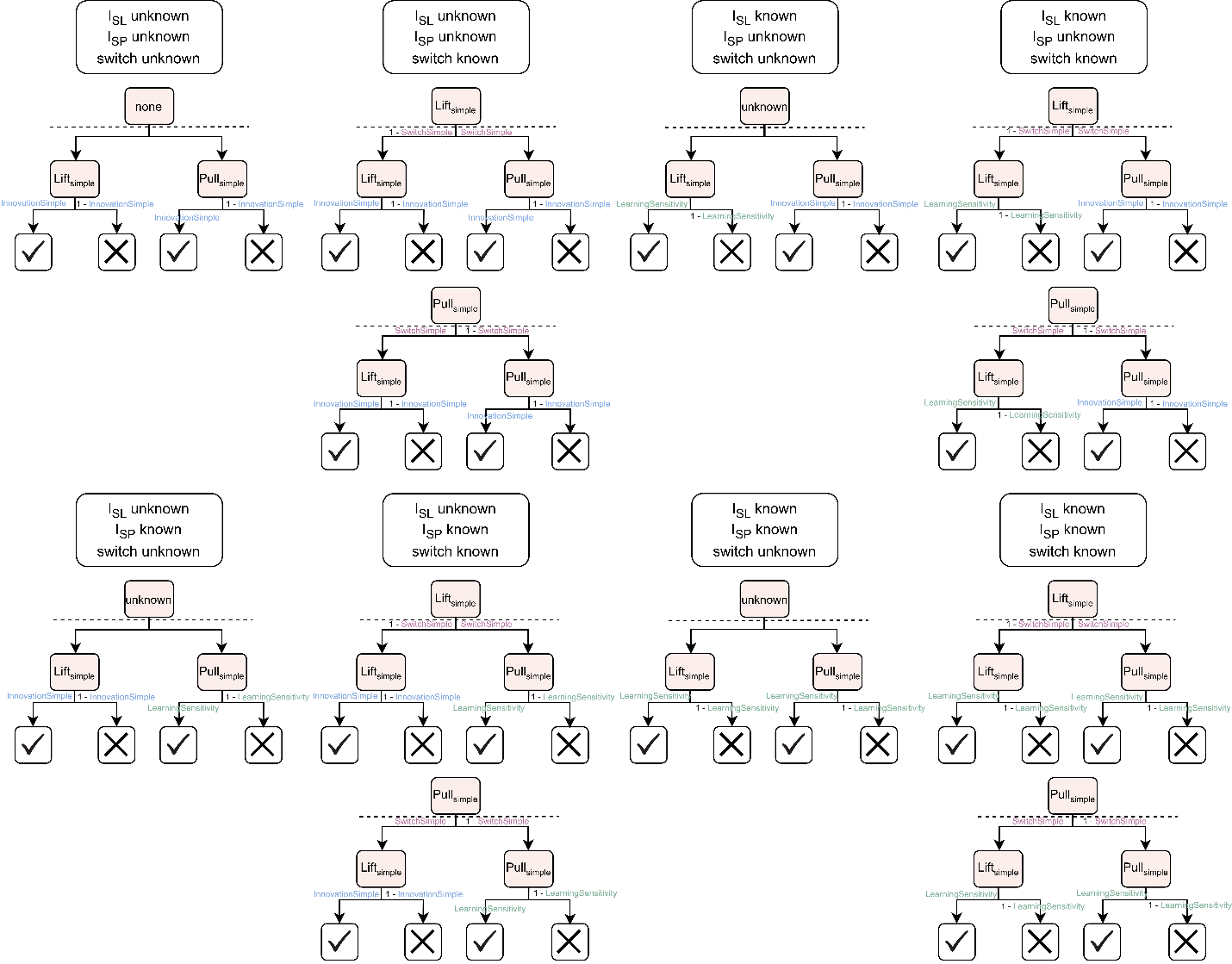

**Phase 2**
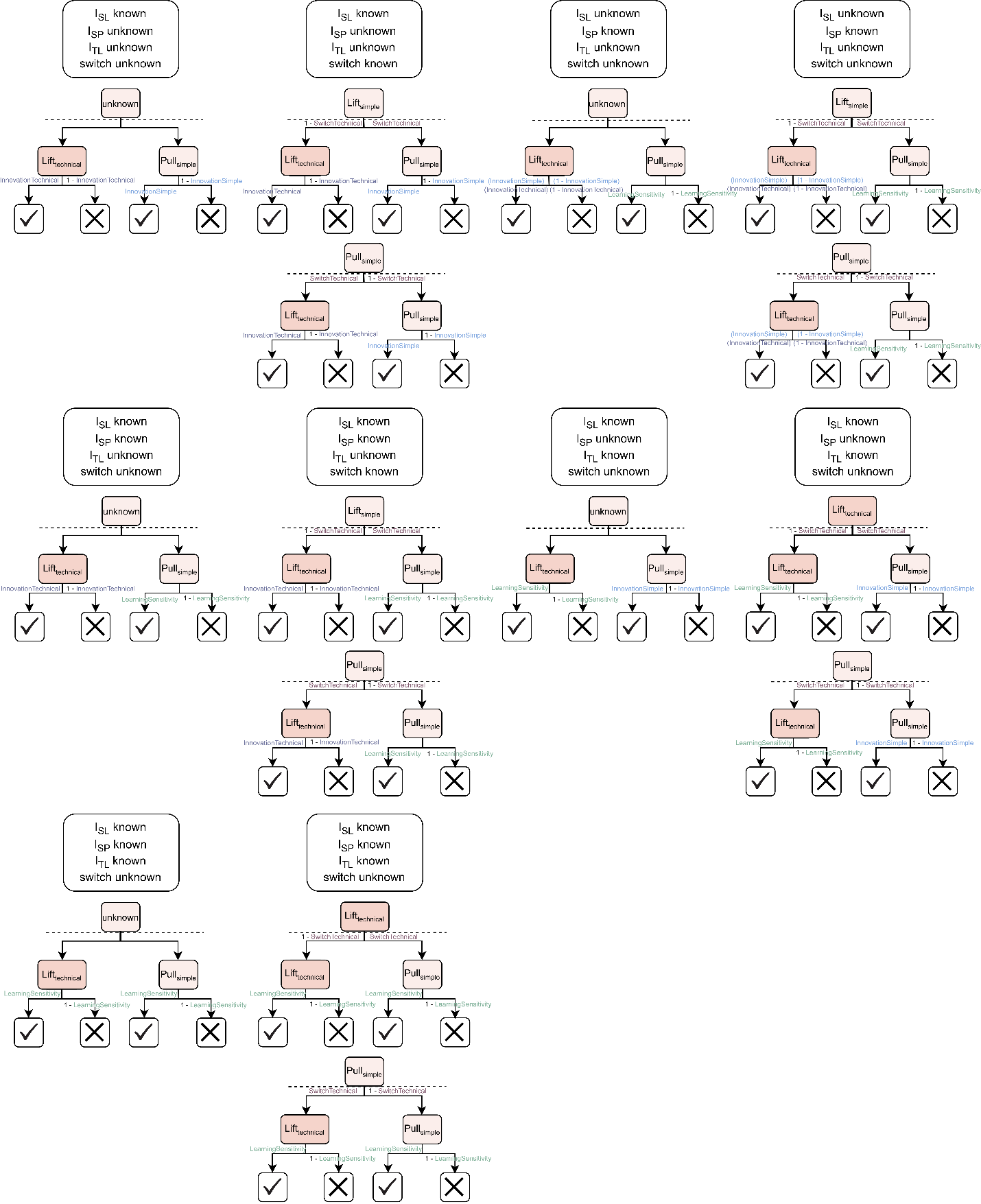

**Phase 3**

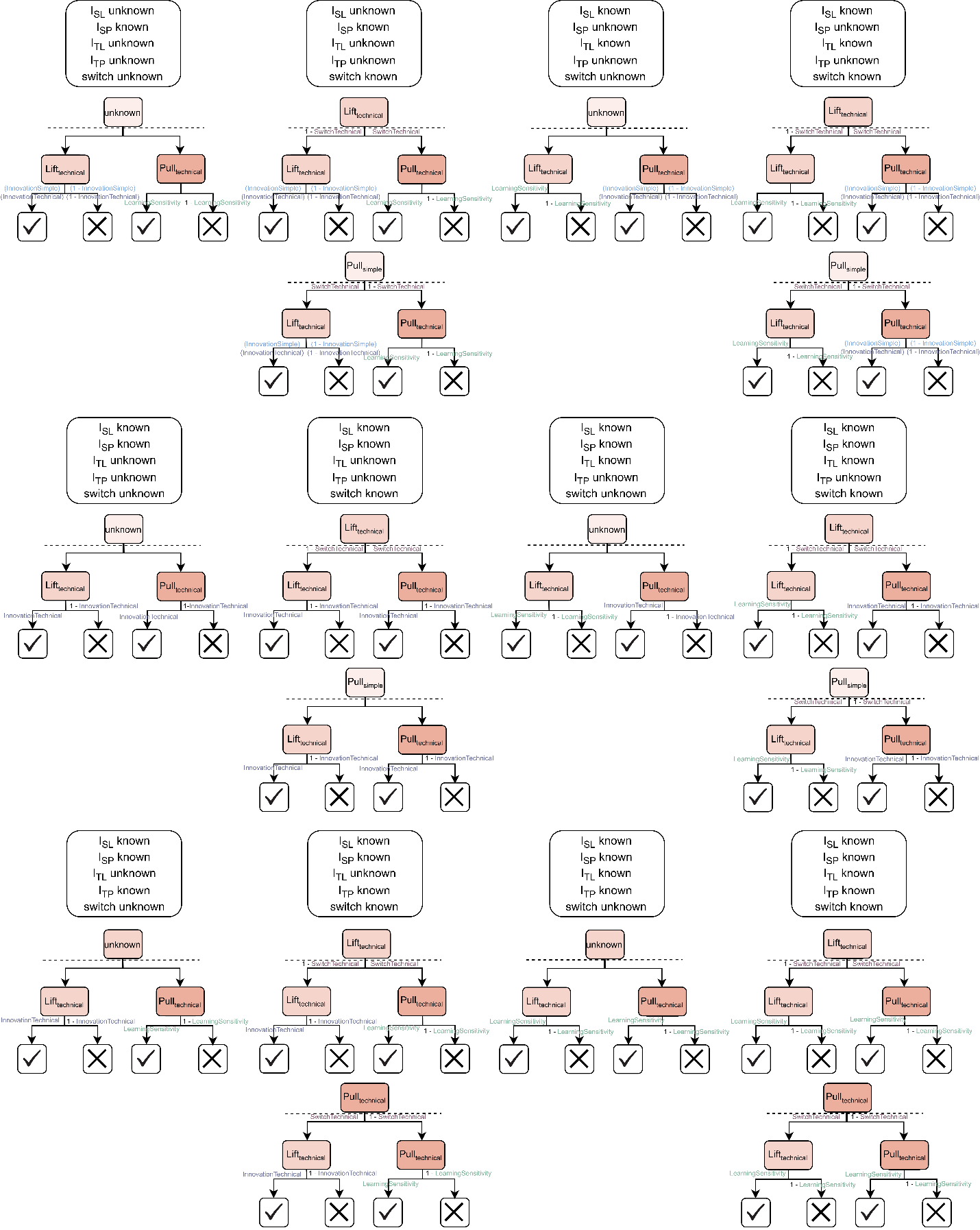

**Figure S2.** Series of decision trees that were used to estimate the probability of observing a given behaviour (e.g., lifting to access the peanut in phase 1, pulling to access the peanut in phase 3). The dashed line delineates the two trials, as the previous trial outcome is used to determine whether the individuals switch in the current trial. Each branch represents a potential pathway (see Methods section). For attempts, both unsuccessful branches are taken. For successes, the respective branch is taken. When the switch is unknown (that is, the previous trial involved attempts on both sides), the switch tendency parameter is not updated. In such cases, only the innovation and/or learning sensitivity parameters are updated for that trial.

*7. Model estimates of cognitive parameters from observed data*

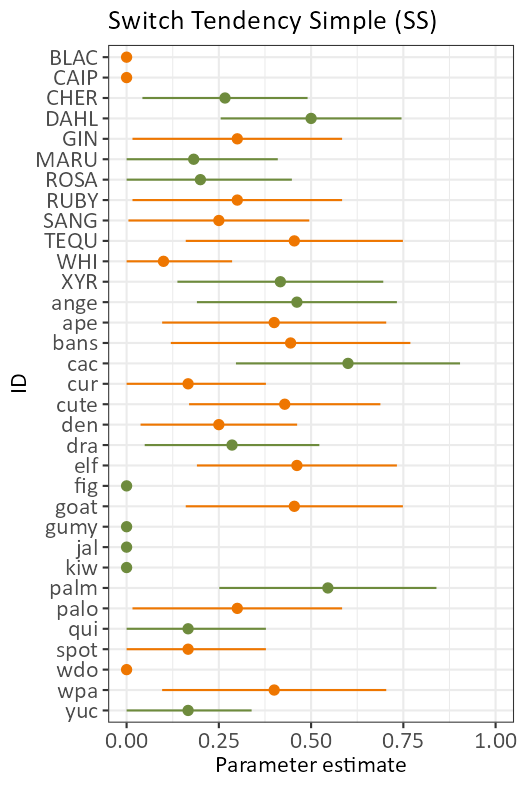

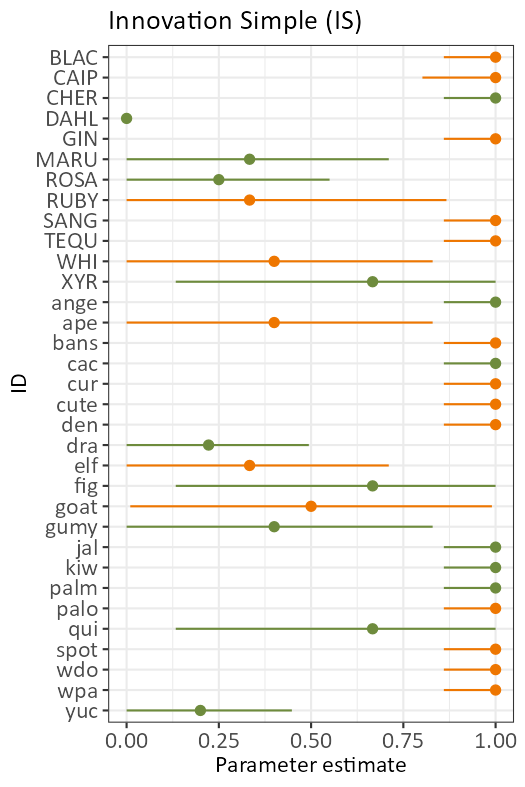

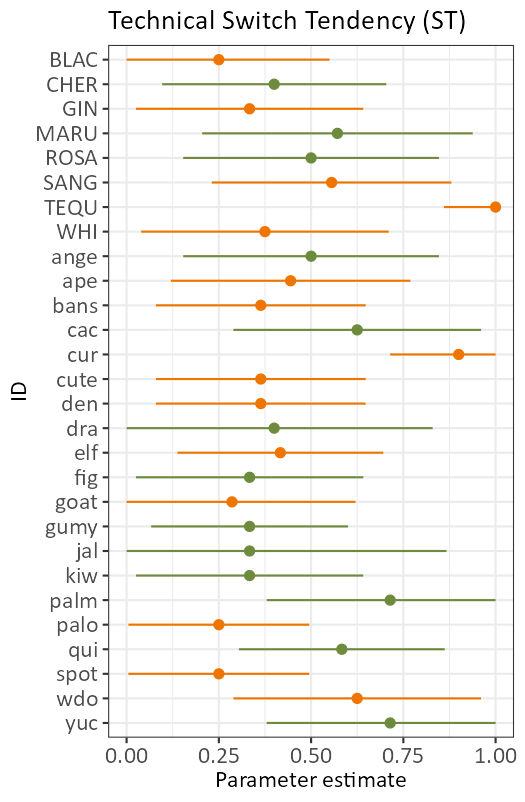

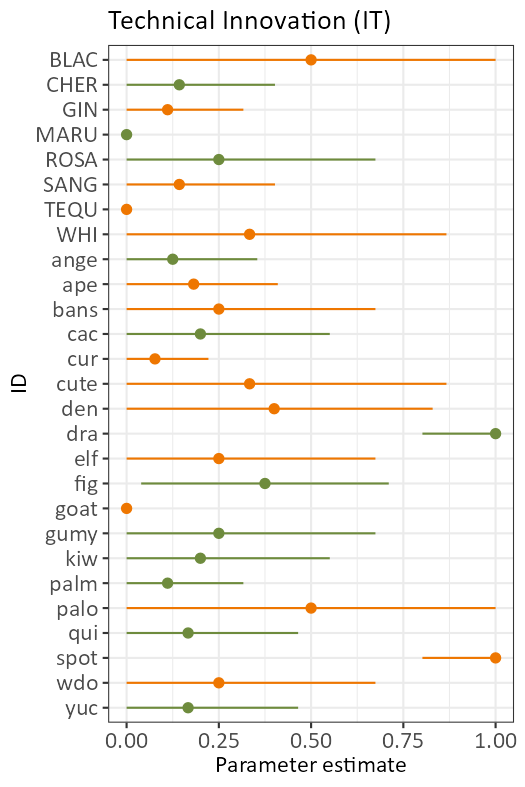

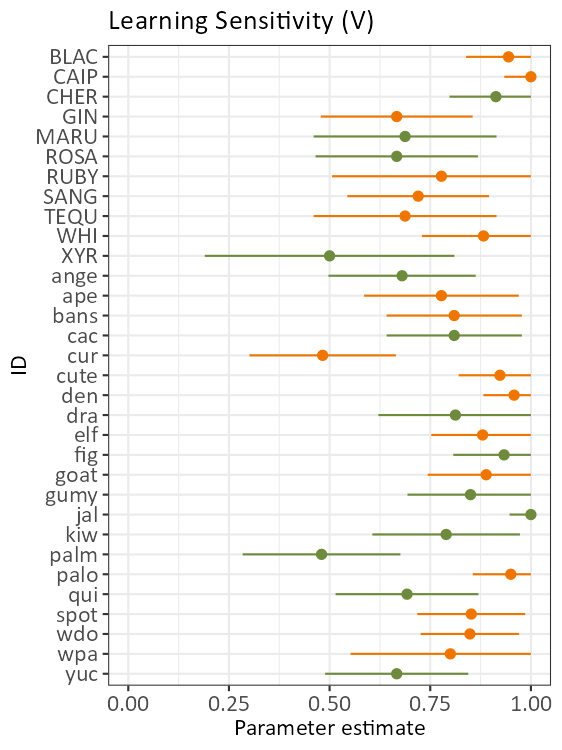

**Figure S3.** Model estimates of parameters from observed data. Each point represents the corresponding evaluated parameter estimate for the specified individual. Higher values indicate better performance for that trait for that individual. The lines indicate the 95% confidence interval for that estimate. For individuals on the *y* axis, capital letters indicate adults while juveniles are in lowercase and four-letter names indicate females, while three-letter names indicate males. Colours indicate troop membership (“Acacia” troop in green and “Savanna” troop in orange).

*8. Quantifying the model inference robustness to describe monkeys’ behavioural flexibility, innovativeness and sensitivity to previous experience*

We proceeded in two steps to quantify the robustness of the inference of the five parameters describing the monkeys’ behaviour in the puzzle-box experiment (i.e., the tendency to switch between known solutions, either simple, *SS*, or technical, *ST*; the capacity to innovate new solutions not performed before during the experiment, either simple, *IS*, or technical, *IT*; the capacity to reproduce known solutions, *V*).

*8.1 Phase length verification*

First, we investigated *a posteriori* whether the phase lengths were sufficient to have reproducible results. To do so, we simulated trial sequences when all parameters were fixed to 0.5. First, we considered only phase 1 and 2 (varying their number of trials between 1 to 15, and 1 to 10, respectively). Next, we considered the number of *successes* *required* on phase 1 before moving to phase 2, which we varied between 1 to 15 too (although capped this at 22 trials as that was the maximum number of trials we actually tested). Third, we varied the number of trials of phase 2 and 3 between 1 to 10. The range of trials were chosen to encapsulate phase lengths below and above the number of trials chosen for our study. For this test, we set each parameter to 0.5 and an initial trial of lifting, pulling, or only attempting (on both sides), we simulated a sequence of trials. We then calculated the parameters from this simulated sequence and repeated it 150 times. By checking the error (i.e., how far each simulated parameter differed from the initial value of 0.5), we checked the quality of the mechanistic model. Overall, the three parameters of main interest, *SS*, *IT*, and *V*, were robustly estimated for at least 10 trials in phase 1, and 4 trials in phase 2, respectively (figure S4, figure S5). Considering phase 3 however highlighted that, given our sampling effort, our estimations are likely to *underestimate* the true parameter values (figure S6).

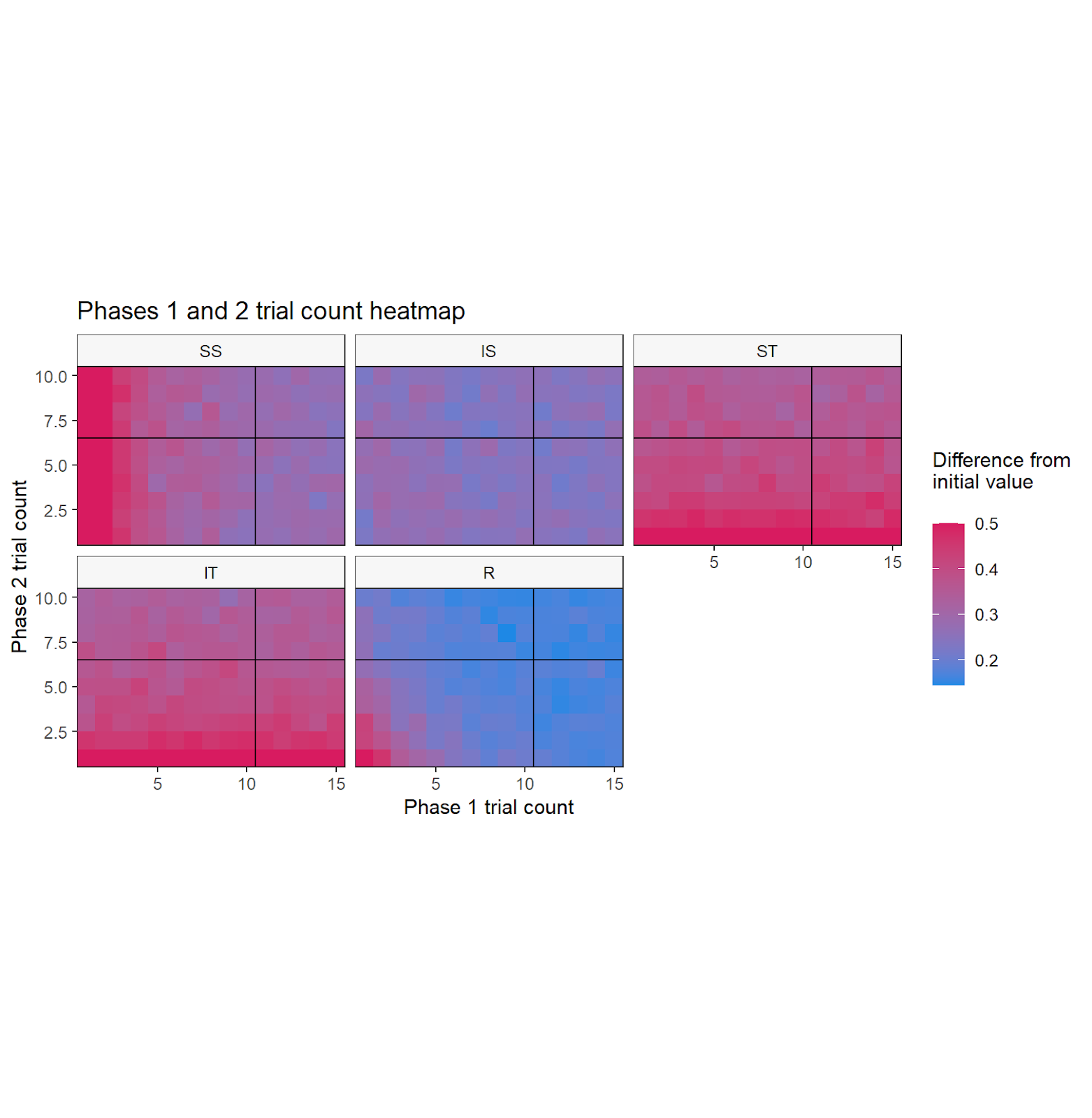

V

**Figure S4**. The mean of each parameter’s difference from the initial value of 0.5 at each combination of phase lengths. Parameters are simulated from 150 repetitions of simulated trial sequences. Bluer areas indicate lower errors for that combination of phase lengths, redder areas indicate higher errors. The solid lines indicate the number of trials used in this experiment. The axes indicate the number of trials in phase 1 and 2. The error for *SS* and *V* improved with the phase 1 trial count. The error for *ST*, *IT*, and *V* improved with the phase 2 trial counts. We conclude that the chosen number of trials is appropriate for each phase.

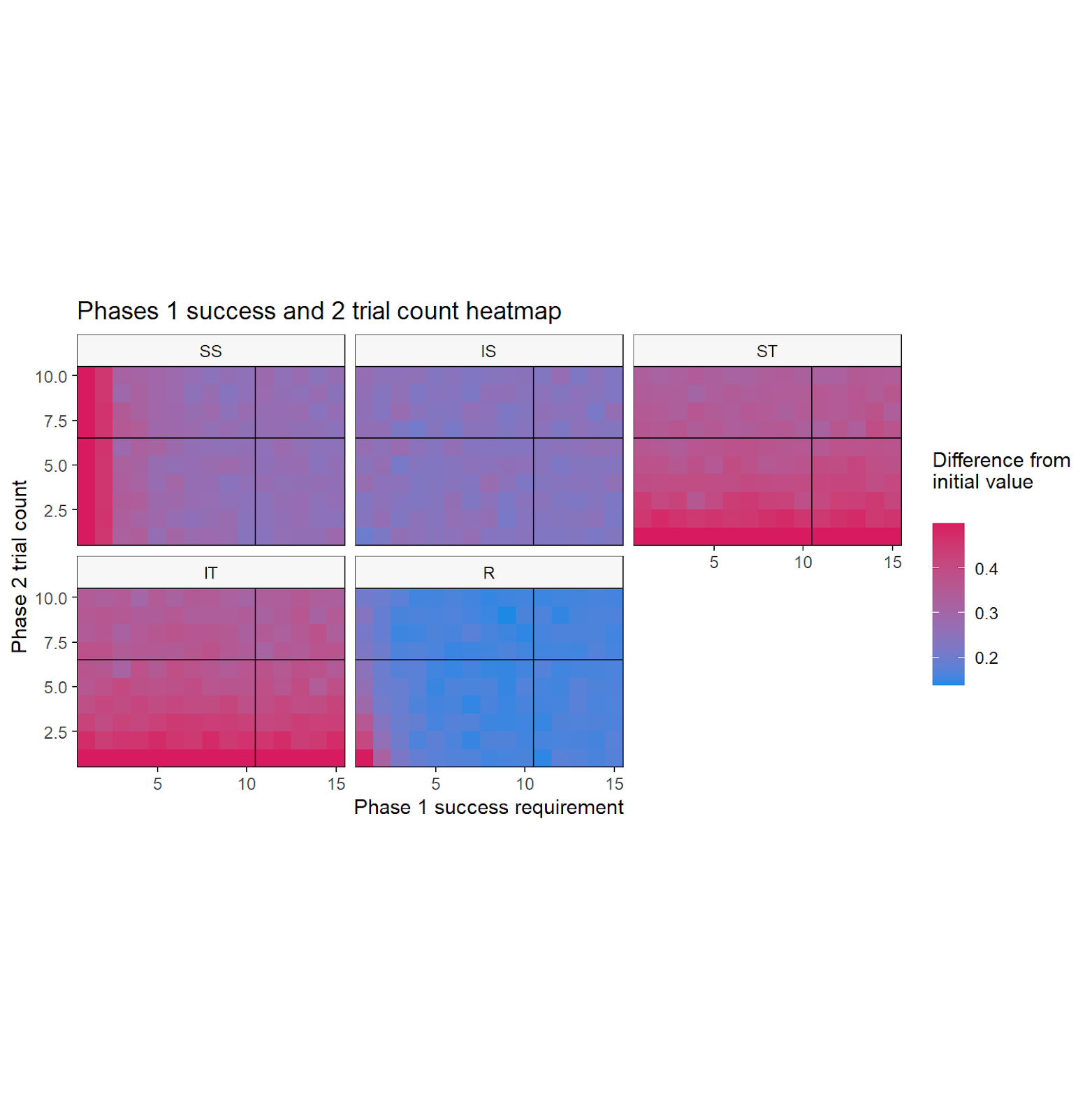

V

**Figure S5**. The mean of each parameter’s difference from the initial value of 0.5 at each combination of phase lengths. Parameters are simulated from 150 repetitions of simulated trial sequences. Bluer areas indicate lower errors, redder areas indicate higher errors. The solid lines indicate the number of trials used in this experiment. The error for *SS* and *V* improved with the phase 1 success count. The error for *ST*, *IT*, and *V* improved with the phase 2 trial counts. We conclude that the chosen number of trials is appropriate for each phase.

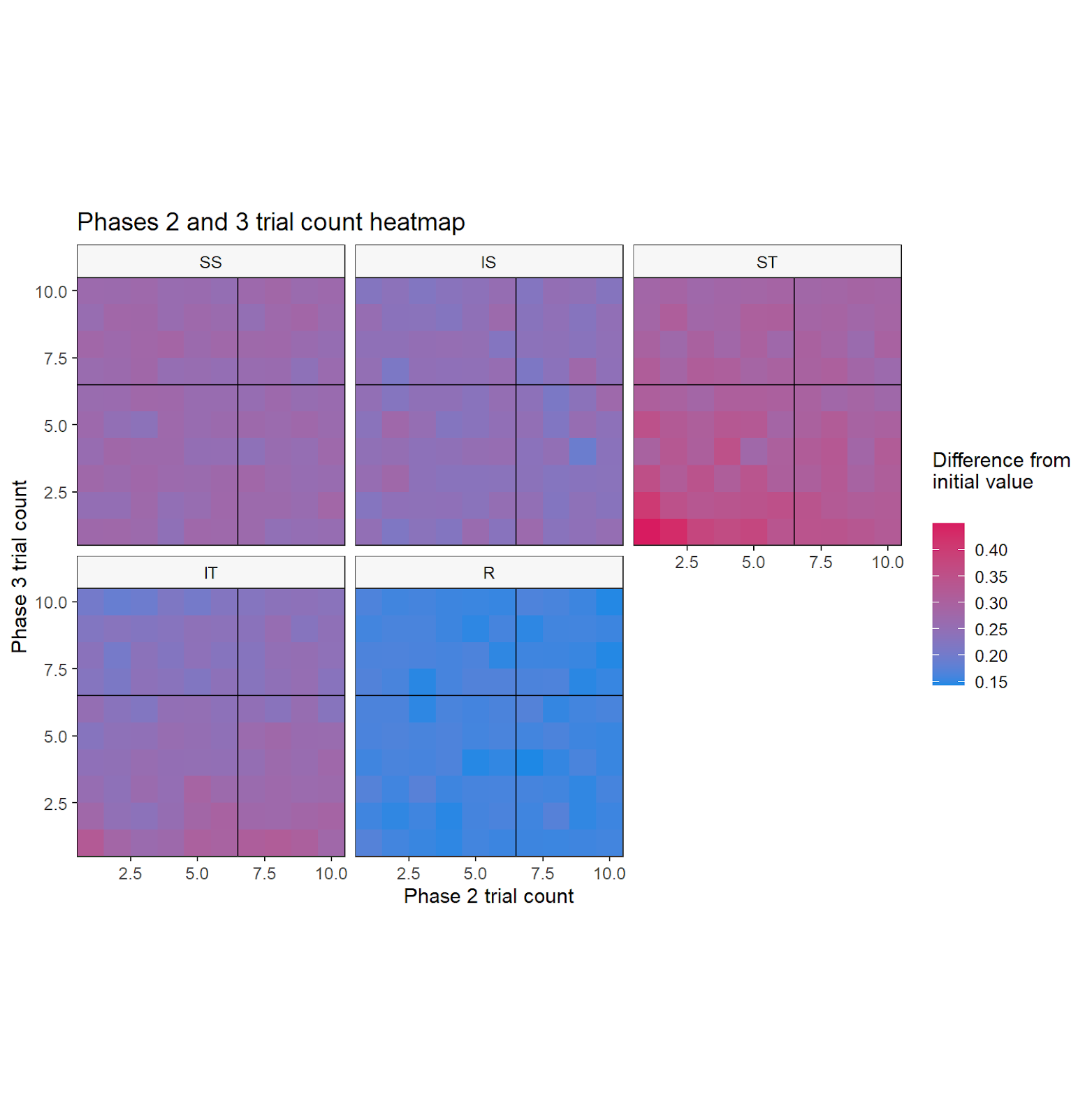

V

**Figure S6**. The mean of each parameter’s difference from the initial value of 0.5 at each combination of phase lengths. Parameters are simulated from 150 repetitions of simulated trial sequences. Bluer areas indicate lower errors, redder areas indicate higher errors. The solid lines indicate the number of trials used in this experiment. The error for *ST* improved with the phase 2 trial count. The error for *ST* and *IT* improved with the phase 2 trial counts. We conclude that the chosen number of trials is appropriate for each phase.

*8.2 Simulated parameter values*

Second, to check the quality of the mechanistic model in determining the parameters, we used *a posteriori* predictive check to see whether we could obtain similar estimations when new data were simulated based on the empirically estimated parameter values (Gelman et al. 1996). To do so, for each individual, we used the estimated parameters to recreate 100 hypothetical trial series. The number of simulated trials matched the original criteria for phase 1, where they passed after 10 successes. The number of trials for phases 2 and 3 matched the observed total trial counts for that vervet monkey. To verify whether the parameters estimated from these hypothetical trials matched those empirically estimated, we refitted the probabilistic model (see main text 2.4.1). That is, we used the decision trees to determine the parameters from the trial sequence. We then compared the “empirical” estimation (based on the experiment) and the “simulated” estimations using on this backward procedure (figures S7 and S8). Overall, our estimations were robust, slightly overestimating the parameter values as a whole (average difference range in parameter values, the reference being the empirical estimate = [0.007, 0.262]; figure S7), but this varied within individuals (figure S8).

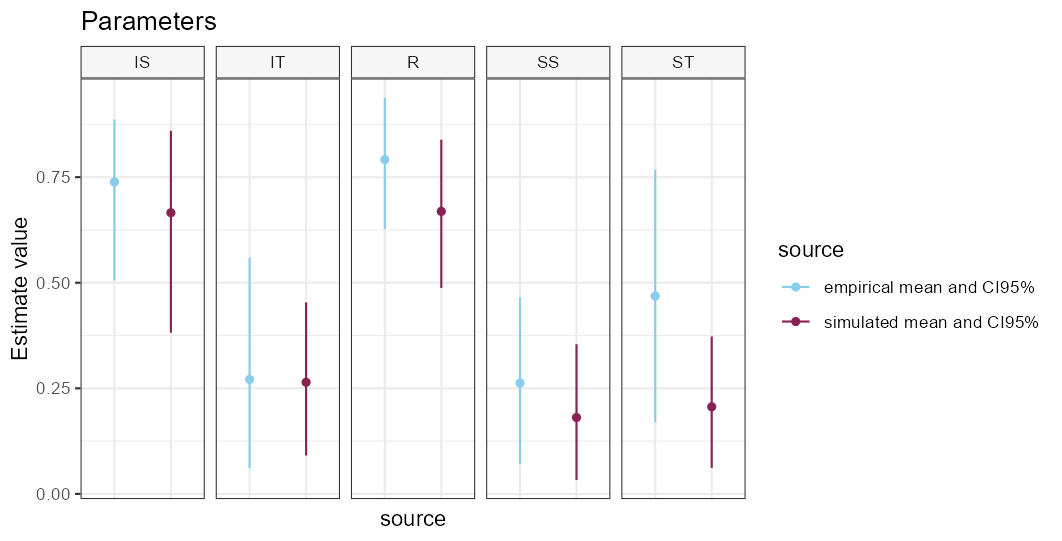

V

**Figure S7.** Robustness of empirically estimated parameters characterising the monkeys’ problem-solving behaviour (populational level). The graphics show the mean of each parameter across all vervet monkeys based on the empirically derived parameters from the observed trial outcomes, compared to the simulated means (100 simulations for each monkey set of parameters). The segments depict the 95% confidence intervals. We conclude that all the parameters are close to the original empirical estimates.

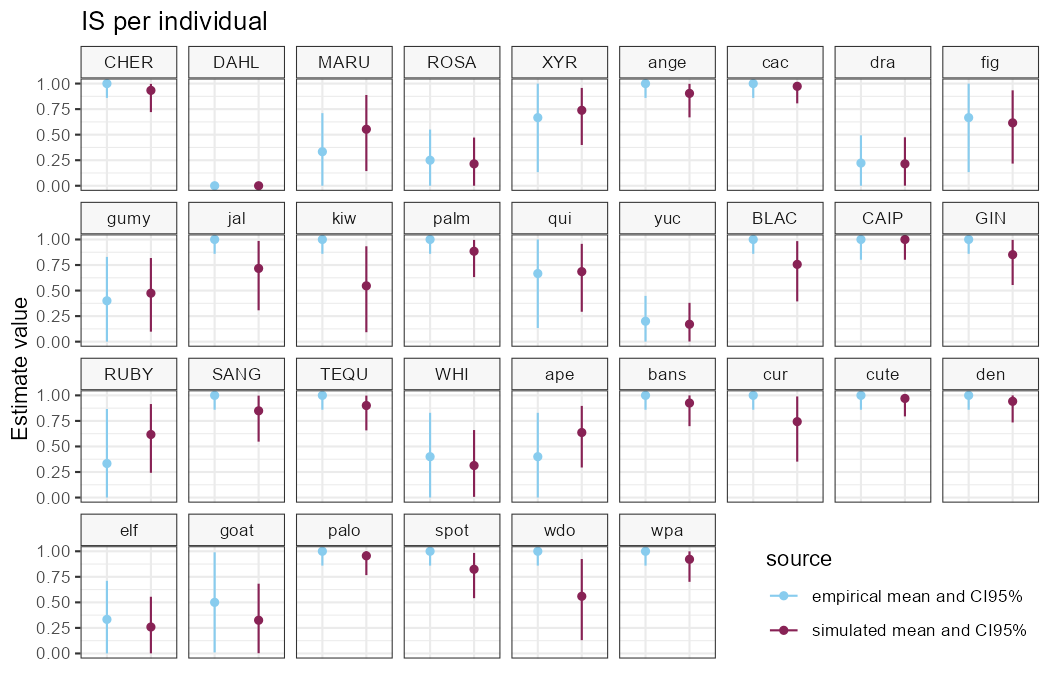

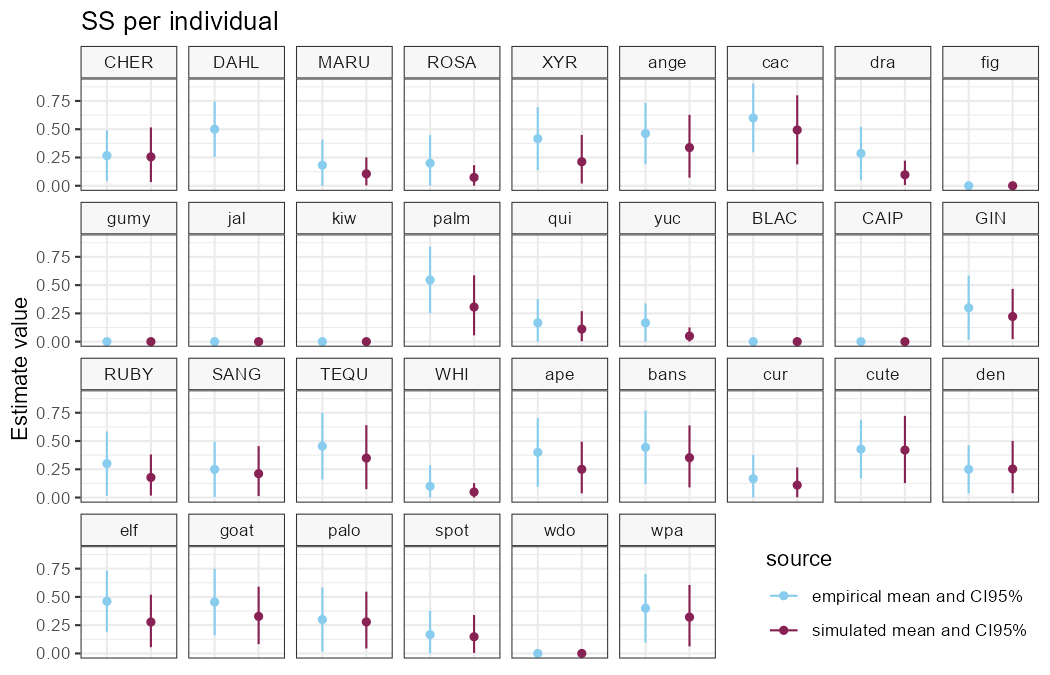

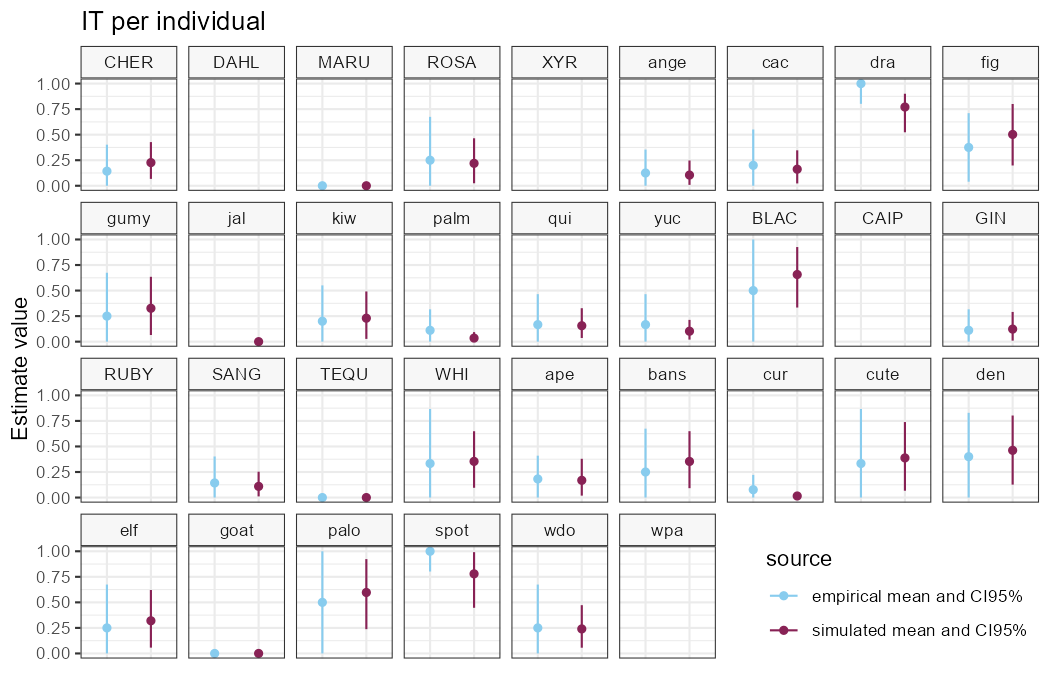

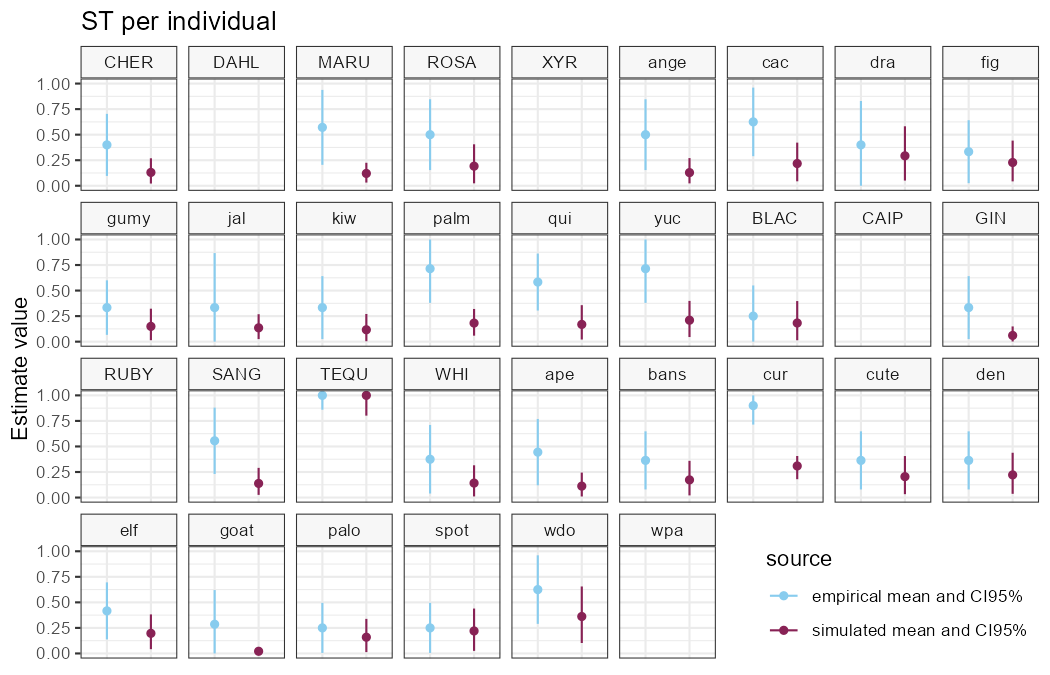

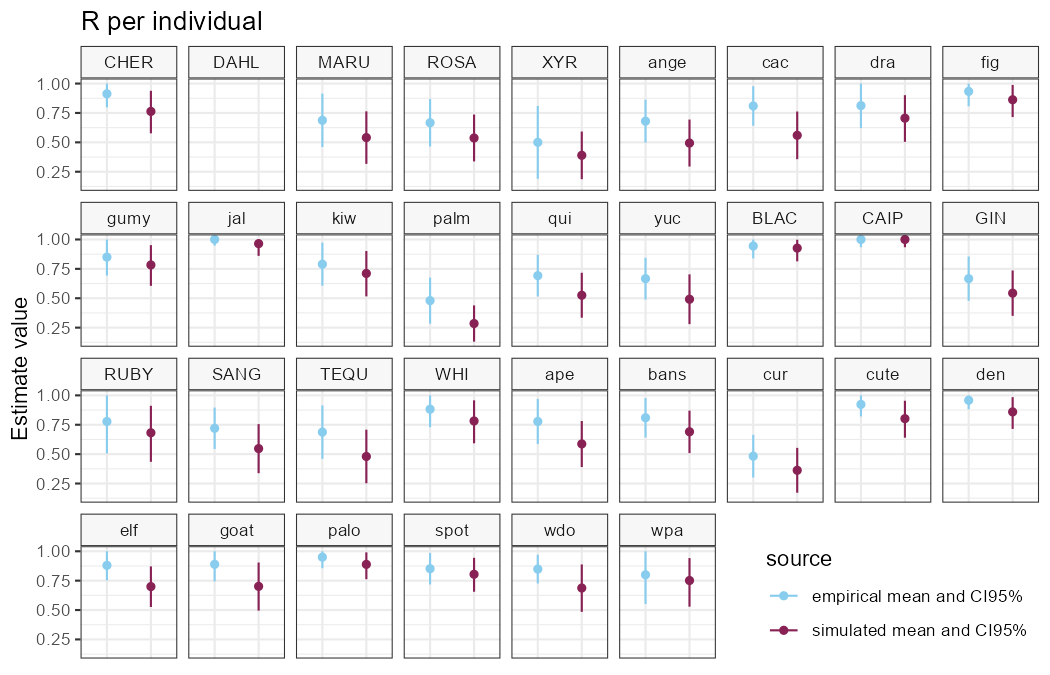

V

**Figure S8**. Robustness of empirically estimated parameters characterising the monkeys’ problem-solving behaviour (individual level). The graphics show the empirically derived parameters from the observed trial outcomes compared to the simulated parameter score for each parameter and for each vervet monkey (100 simulations for each monkey set of parameters). Each facet plot indicates one individual, where capital letters indicate an adult while juveniles are written in lowercase, and four-letter names indicate females, while three-letter names indicate males. The first set of adults and juveniles is from the first troop (“Acacia”) and the second set is from the second troop (“Savanna”). Overall, the simulated parameters are close to the empirical ones.

*9. Principal Component Analysis: correlation check and visualization*

Our analyses considered the global effects of anthropogenic raiding experience, demographic traits, and previous social exposure to the box. The composite demographic and social variables were obtained by considering the first component of a Principal Component Analysis (PCA, “prcomp” function of the *stats* package) summarizing age, sex and rank (demographic variables; the first axis representing 45.8% of the variance; figure S9; rank determination in Supp. Mat. S10; Elo, 1978; Neumann & Kulik, 2020), and the number of trials with the same solution that the monkey first used prior to their first opening on phase 1 (reflecting exposure to opening techniques), the number of innovation trials the individual viewed prior to starting phase 2 (reflecting exposure to technical innovation), and the number of phase 2 trials the individual saw within 5 m prior to starting phase 2 (reflecting exposure to the experimental challenge overall); the first axis representing 42.04% of the variance; figure S10). Prior to performing the PCAs, we scaled all continuous variables (to a mean of 0 and a standard deviation of 1) to ensure that all variables have the same influence and verified that variables were lowly correlated with each other (Spearman’s correlation < 0.6, figures S9 and S10). For the social variables, the counts were all restricted to two-week time periods, as we considered that this is a salient time period for monkey memory (Bakner & Treichler, 1993 but see Supp. Mat. S11 for a sensitivity analysis when using smaller/larger time thresholds).

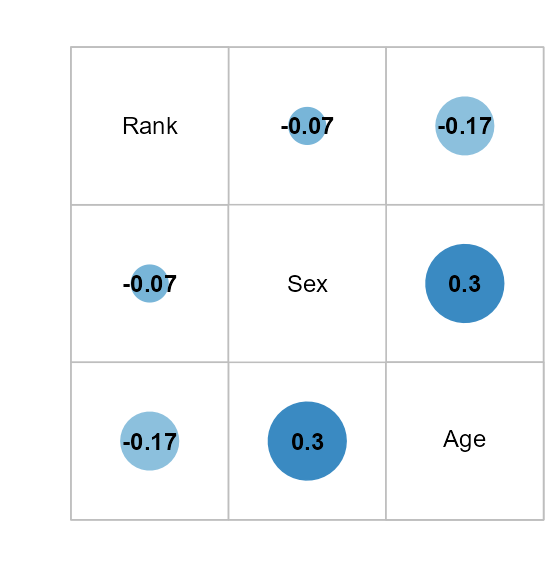

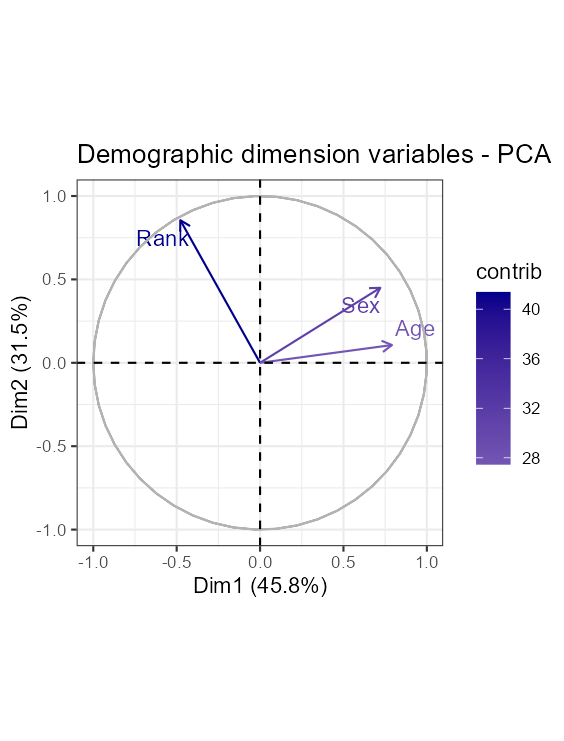

**Figure S9**. Demographic dimension correlogram (left) where the size of the points is a function of the correlation (Pearson correlation; “rcorr” function of the *Hmisc* package, Harrell, 2019). PCA output (right) where the arrows indicate the loads of each variable and projected scaled data, mapped into a unit circle (“fviz_pca_var” function of the *factoextra* package, Kassambara, 2020). Contrib = contribution (percentage of variance explained). We used dimension 1 as the global demographic variable in analyses.

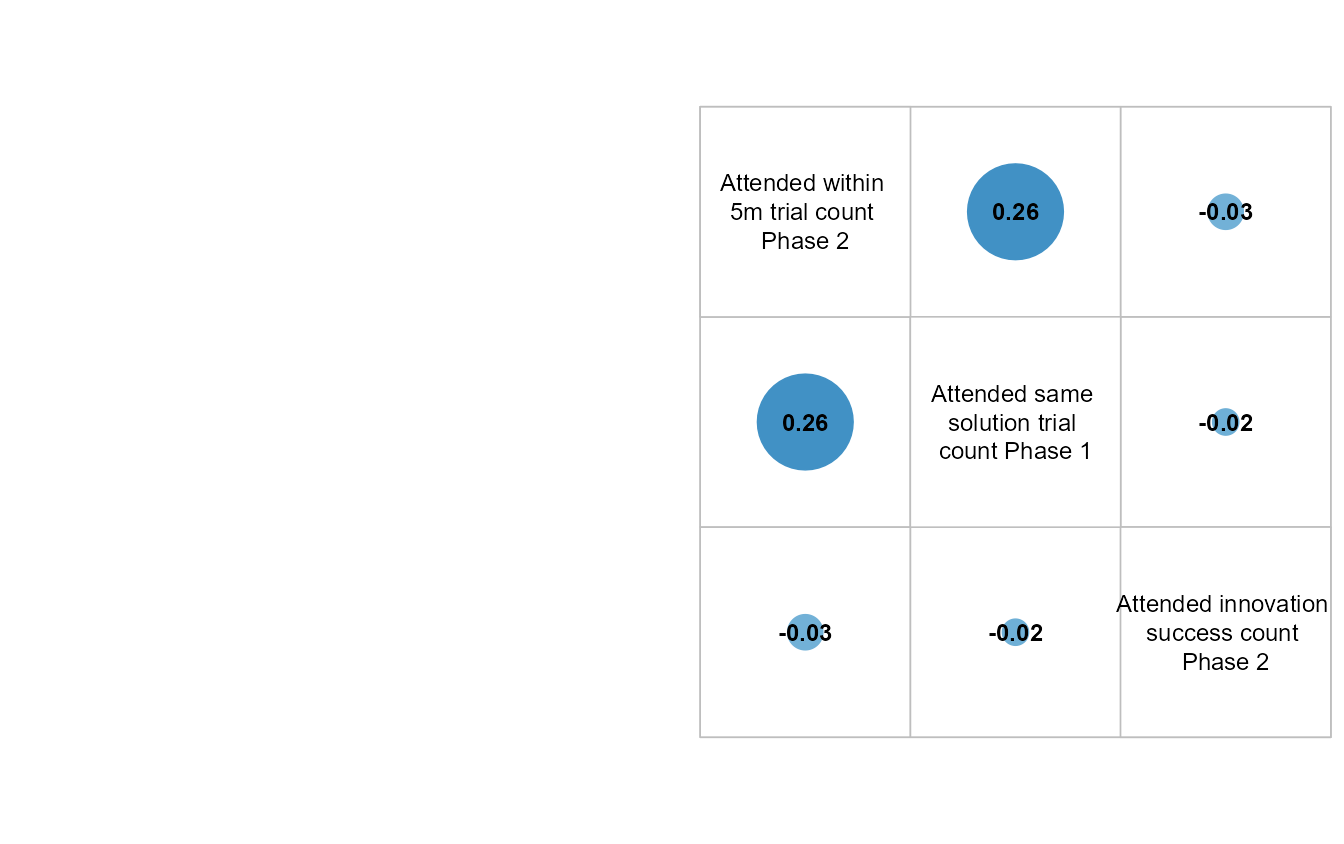

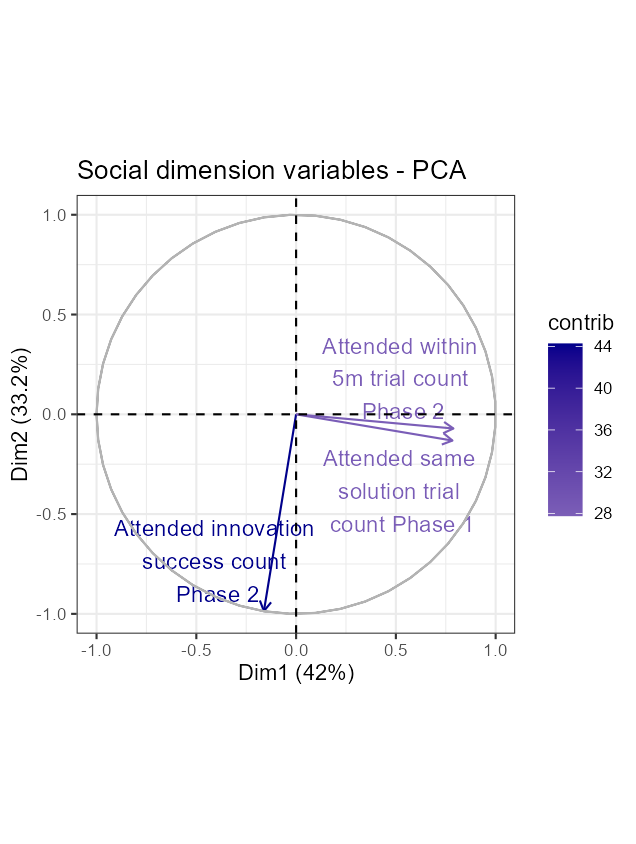

**Figure S10**. Social dimension correlogram (left) where the size of the points is a function of the correlation (Pearson correlation; “rcorr” function of the *Hmisc* package, Harrell, 2019). PCA output (right) where the arrows indicate the loads of each variable and projected scaled data, mapped into a unit circle (“fviz_pca_var” function of the *factoextra* package, Kassambara, 2020). Contrib = contribution (percentage of variance explained). We used dimension 1 as the global social variable in analyses.

*10. Rank determination*

We used all dyadic social agonistic observations recorded between September 2023 and February 2024 (ethogram Supp. Mat. table S3; 210 interactions in the two troops), to estimate an individuals’ social rank within their troop. We determined individual hierarchical ranks using the Elo-scores method (Elo, 1978) using the “elo.seq” function of the *EloRating* package, accounting for temporal changes in social dynamics over time (Neumann & Kulik, 2020). An actor ‘won’ the interaction when the victim displayed submissive behaviour such as fleeing, crawling, or retreating, thus ending and ‘losing’ the interaction. We averaged the scores per individual over the study period to assess their average rank while participating in the study. Scores were then standardized to a mean of 0 and standard deviation of 1 within their troop.

*11. Analyses’ sensitivity to arbitrary decisions*

Our analyses relied on two relatively arbitrary choices: the threshold to consider the time threshold to measure prior exposure to the task (fixed to a two-week period in the main analyses) and whether an individual is a technical innovator or not (with regards to the technical innovation term, *IT*, comprised between 0 and 1; fixed to 0.5 in the main analyses). To explore the impact of these choices, we varied the threshold of social exposure to the experiment in the SS and ‘good IT’ models from 7 to 21 days. In addition, we varied the ‘good IT’ cutoff threshold from the upper limit of the CI_95%_ from 0.3 to 0.7 (tables S3 and S4; figures S10 and S11).

Across all variations, technical innovation is explained more by demographic traits and previous social exposure to the experiment factors, but not anthropogenic raiding experience, which highlights little sensitivity of our inference to any of these two choices (table S5 and figure S11, and table S6 and figure S12 for sensitivity to the social exposure time window threshold, and the good *IT* threshold, respectively).

*11.1 SS model: days of social exposure to box sensitivity*

**Table S5**. Comparison of models investigating the effects of the anthropogenic foraging experience, demographic, and social exposure to the box dimensions on simple switch tendency scores, where the social exposure to the experiment varies. Here, we tested the sensitivity of this period by varying the number of days from 7 to 21. *K* represents the number of estimated parameters for each model, *AICc* the Akaike Information Criterion corrected for small samples, *∆AIC* is obtained by comparing to the most parsimonious model (first row), log-likelihood is the probability of the observed data, *weight* corresponds to the Akaike weights (exp(-0.5**AICc*)), and their cumulative weight.

| *7 days* | *Model names* | *K* | *AICc* | *ΔAICc* | *Log-likelihood* | *Weight* | *Cumulative weight* | |
| --- | --- | --- | --- | --- | --- | --- | --- | --- |
|  | ***null*** | **2** | **-22.61** | **0.00** | **13.51** | **0.64** | | **0.64** |
|  | **Experience** | **3** | **-20.96** | **1.65** | **13.90** | **0.28** | | **0.92** |
|  | **Demographic** | **3** | **-16.09** | **6.52** | **11.57** | **0.02** | | **0.95** |
|  | Social | 3 | -15.97 | 6.64 | 11.51 | 0.02 | | 0.97 |
|  | Social, Experience | 4 | -14.15 | 8.46 | 11.99 | 0.01 | | 0.98 |
|  | Demographic, Experience | 4 | -14.09 | 8.52 | 11.95 | 0.01 | | 0.99 |
|  | Demographic, Social | 4 | -13.67 | 8.94 | 11.75 | 0.01 | | 1.00 |
|  | Demographic, Social, Experience | 5 | -11.23 | 11.38 | 12.04 | 0.00 | | 1.00 |

| *10 days* | *Model names* | *K* | *AICc* | *ΔAICc* | *Log-likelihood* | *Weight* | *Cumulative weight* | |
| --- | --- | --- | --- | --- | --- | --- | --- | --- |
|  | ***null*** | **2** | **-22.61** | **0.00** | **13.51** | **0.61** | | **0.61** |
|  | **Experience** | **3** | **-20.96** | **1.65** | **13.90** | **0.27** | | **0.88** |
|  | **Social** | **3** | **-17.32** | **5.29** | **12.18** | **0.04** | | **0.93** |
|  | **Demographic** | **3** | **-16.09** | **6.52** | **11.57** | **0.02** | | **0.95** |
|  | Social, Experience | 4 | -15.72 | 6.90 | 12.77 | 0.02 | | 0.97 |
|  | Demographic, Social | 4 | -15.39 | 7.22 | 12.60 | 0.02 | | 0.99 |
|  | Demographic, Experience | 4 | -14.09 | 8.52 | 11.95 | 0.01 | | 1.00 |
|  | Demographic, Social, Experience | 5 | -12.95 | 9.66 | 12.90 | 0.00 | | 1.00 |

| *12 days* | *Model names* | *K* | *AICc* | *ΔAICc* | *Log-likelihood* | *Weight* | *Cumulative weight* | |
| --- | --- | --- | --- | --- | --- | --- | --- | --- |
|  | ***null*** | **2** | **-22.61** | **0.00** | **13.51** | **0.64** | | **0.64** |
|  | **Experience** | **3** | **-20.96** | **1.65** | **13.90** | **0.28** | | **0.93** |
|  | **Demographic** | **3** | **-16.09** | **6.52** | **11.57** | **0.02** | | **0.95** |
|  | Social | 3 | -15.68 | 6.93 | 11.36 | 0.02 | | 0.97 |
|  | Demographic, Experience | 4 | -14.09 | 8.52 | 11.95 | 0.01 | | 0.98 |
|  | Social, Experience | 4 | -14.07 | 8.54 | 11.94 | 0.01 | | 0.99 |
|  | Demographic, Social | 4 | -13.57 | 9.04 | 11.69 | 0.01 | | 1.00 |
|  | Demographic, Social, Experience | 5 | -11.19 | 11.42 | 12.02 | 0.00 | | 1.00 |

| *14 days* | *Model names* | *K* | *AICc* | *ΔAICc* | *Log-likelihood* | *Weight* | *Cumulative weight* | |
| --- | --- | --- | --- | --- | --- | --- | --- | --- |
|  | ***null*** | **2** | **-22.61** | **0.00** | **13.51** | **0.64** | | **0.64** |
|  | **Experience** | **3** | **-20.96** | **1.65** | **13.90** | **0.28** | | **0.92** |
|  | **Demographic** | **3** | **-16.09** | **6.52** | **11.57** | **0.02** | | **0.95** |
|  | Social | 3 | -15.90 | 6.71 | 11.47 | 0.02 | | 0.97 |
|  | Social, Experience | 4 | -14.43 | 8.18 | 12.13 | 0.01 | | 0.98 |
|  | Demographic, Experience | 4 | -14.09 | 8.52 | 11.95 | 0.01 | | 0.99 |
|  | Demographic, Social | 4 | -13.82 | 8.79 | 11.82 | 0.01 | | 1.00 |
|  | Demographic, Social, Experience | 5 | -11.56 | 11.05 | 12.21 | 0.00 | | 1.00 |

| *16 days* | *Model names* | *K* | *AICc* | *ΔAICc* | *Log-likelihood* | *Weight* | *Cumulative weight* | |
| --- | --- | --- | --- | --- | --- | --- | --- | --- |
|  | ***null*** | **2** | **-22.61** | **0.00** | **13.51** | **0.64** | | **0.64** |
|  | **Experience** | **3** | **-20.96** | **1.65** | **13.90** | **0.28** | | **0.92** |
|  | **Demographic** | **3** | **-16.09** | **6.52** | **11.57** | **0.02** | | **0.95** |
|  | Social | 3 | -16.02 | 6.59 | 11.53 | 0.02 | | 0.97 |
|  | Social, Experience | 4 | -14.50 | 8.12 | 12.16 | 0.01 | | 0.98 |
|  | Demographic, Experience | 4 | -14.09 | 8.52 | 11.95 | 0.01 | | 0.99 |
|  | Demographic, Social | 4 | -13.95 | 8.66 | 11.88 | 0.01 | | 1.00 |
|  | Demographic, Social, Experience | 5 | -11.64 | 10.97 | 12.25 | 0.00 | | 1.00 |

| *18 days* | *Model names* | *K* | *AICc* | *ΔAICc* | *Log-likelihood* | *Weight* | *Cumulative weight* | |
| --- | --- | --- | --- | --- | --- | --- | --- | --- |
|  | ***null*** | **2** | **-22.61** | **0.00** | **13.51** | **0.64** | | **0.64** |
|  | **Experience** | **3** | **-20.96** | **1.65** | **13.90** | **0.28** | | **0.92** |
|  | **Social** | **3** | **-16.21** | **6.40** | **11.63** | **0.03** | | **0.94** |
|  | **Demographic** | **3** | **-16.09** | **6.52** | **11.57** | **0.02** | | **0.97** |
|  | Social, Experience | 4 | -14.72 | 7.89 | 12.27 | 0.01 | | 0.98 |
|  | Demographic, Social | 4 | -14.20 | 8.41 | 12.01 | 0.01 | | 0.99 |
|  | Demographic, Experience | 4 | -14.09 | 8.52 | 11.95 | 0.01 | | 1.00 |
|  | Demographic, Social, Experience | 5 | -11.89 | 10.72 | 12.38 | 0.00 | | 1.00 |

| *21 days* | *Model names* | *K* | *AICc* | *ΔAICc* | *Log-likelihood* | *Weight* | *Cumulative weight* | |
| --- | --- | --- | --- | --- | --- | --- | --- | --- |
|  | ***null*** | **2** | **-22.61** | **0.00** | **13.51** | **0.63** | | **0.63** |
|  | **Experience** | **3** | **-20.96** | **1.65** | **13.90** | **0.28** | | **0.91** |
|  | **Social** | **3** | **-16.29** | **6.32** | **11.67** | **0.03** | | **0.94** |
|  | **Demographic** | **3** | **-16.09** | **6.52** | **11.57** | **0.02** | | **0.96** |
|  | Social, Experience | 4 | -15.14 | 7.47 | 12.48 | 0.02 | | 0.98 |
|  | Demographic, Social | 4 | -14.65 | 7.96 | 12.23 | 0.01 | | 0.99 |
|  | Demographic, Experience | 4 | -14.09 | 8.52 | 11.95 | 0.01 | | 1.00 |
|  | Demographic, Social, Experience | 5 | -12.50 | 10.11 | 12.68 | 0.00 | | 1.00 |

 
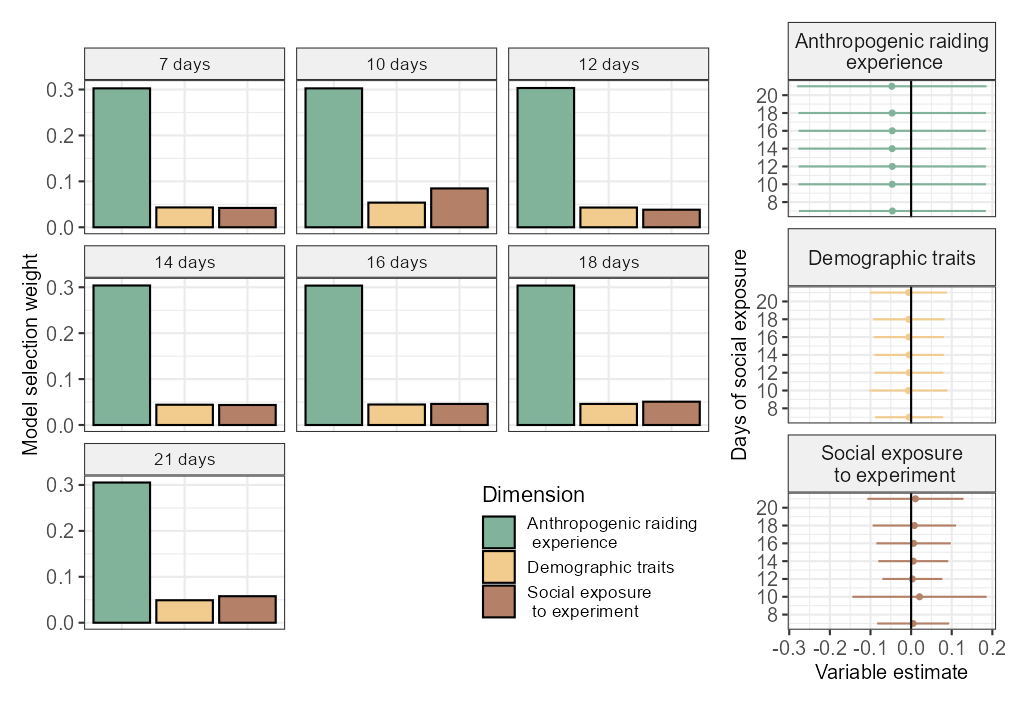

**Figure S11**. Comparison of models investigating effects of the anthropogenic raiding experience, demographic, and social exposure to the box dimensions on simple switch tendency scores, where the social exposure to the experiment varies. In the original analysis, we used a 14-day period (in green) of social exposure to the experiment. Here, we tested the sensitivity of this period by varying the number of days from 7 to 21.

*11.2 ‘good IT’ model: days of social exposure to box sensitivity*

**Table S6**. Comparison of models investigating the effects of the anthropogenic foraging experience, demographic, and social exposure to the box dimensions on good technical innovation scores, where the social exposure to the experiment varies. Here, we tested the sensitivity of this period by varying the number of days from 7 to 21. *K* represents the number of estimated parameters for each model, *AICc* the Akaike Information Criterion corrected for small samples, *∆AIC* is obtained by comparing to the most parsimonious model (first row), log-likelihood is the probability of the observed data, *weight* corresponds to the Akaike weights (exp(-0.5**AICc*)), and their cumulative weight.

| *7 days* | *Model names* | *K* | *AICc* | *ΔAICc* | *Log-likelihood* | *Weight* | *Cumulative weight* |
| --- | --- | --- | --- | --- | --- | --- | --- |
|  | **Demographic** | **2** | **39.78** | **0.00** | **-17.64** | **0.35** | **0.35** |
|  | **Social** | **2** | **40.73** | **0.94** | **-18.11** | **0.22** | **0.57** |
|  | **Demographic, Social** | **3** | **40.99** | **1.21** | **-16.97** | **0.19** | **0.77** |
|  | **Demographic, Experience** | **3** | **42.31** | **2.53** | **-17.64** | **0.10** | **0.87** |
|  | **Social, Experience** | **3** | **42.90** | **3.12** | **-17.93** | **0.07** | **0.94** |
|  | **Demographic, Social, Experience** | **4** | **43.75** | **3.96** | **-16.96** | **0.05** | **0.99** |
|  | *null* | 1 | 47.60 | 7.82 | -22.74 | 0.01 | 1.00 |
|  | Experience | 2 | 49.37 | 9.59 | -22.49 | 0.00 | 1.00 |

| *10 days* | *Model names* | *K* | *AICc* | *ΔAICc* | *Log-likelihood* | *Weight* | *Cumulative weight* | |
| --- | --- | --- | --- | --- | --- | --- | --- | --- |
|  | **Demographic** | **2** | **39.78** | **0.00** | **-17.64** | **0.44** | | **0.44** |
|  | **Social** | **2** | **41.43** | **1.64** | **-18.46** | **0.20** | | **0.64** |
|  | **Demographic, Social** | **3** | **42.24** | **2.45** | **-17.60** | **0.13** | | **0.77** |
|  | **Demographic, Experience** | **3** | **42.31** | **2.53** | **-17.64** | **0.13** | | **0.90** |
|  | **Social, Experience** | **3** | **43.82** | **4.03** | **-18.39** | **0.06** | | **0.95** |
|  | Demographic, Social, Experience | 4 | 45.00 | 5.22 | -17.59 | 0.03 | | 0.99 |
|  | *null* | 1 | 47.60 | 7.82 | -22.74 | 0.01 | | 1.00 |
|  | Experience | 2 | 49.37 | 9.59 | -22.49 | 0.00 | | 1.00 |

| *12 days* | *Model names* | *K* | *AICc* | *ΔAICc* | *Log-likelihood* | *Weight* | *Cumulative weight* | |
| --- | --- | --- | --- | --- | --- | --- | --- | --- |
|  | **Demographic** | **2** | **39.78** | **0.00** | **-17.64** | **0.36** | | **0.36** |
|  | **Social** | **2** | **40.58** | **0.79** | **-18.04** | **0.24** | | **0.61** |
|  | **Demographic, Social** | **3** | **41.43** | **1.65** | **-17.19** | **0.16** | | **0.77** |
|  | **Demographic, Experience** | **3** | **42.31** | **2.53** | **-17.64** | **0.10** | | **0.87** |
|  | **Social, Experience** | **3** | **42.85** | **3.06** | **-17.90** | **0.08** | | **0.95** |
|  | Demographic, Social, Experience | 4 | 44.20 | 4.42 | -17.19 | 0.04 | | 0.99 |
|  | *null* | 1 | 47.60 | 7.82 | -22.74 | 0.01 | | 1.00 |
|  | Experience | 2 | 49.37 | 9.59 | -22.49 | 0.00 | | 1.00 |

| *14 days* | *Model names* | *K* | *AICc* | *ΔAICc* | *Log-likelihood* | *Weight* | *Cumulative weight* | |
| --- | --- | --- | --- | --- | --- | --- | --- | --- |
|  | **Demographic** | **2** | **39.78** | **0.00** | **-17.64** | **0.42** | | **0.42** |
|  | **Social** | **2** | **41.14** | **1.35** | **-18.32** | **0.21** | | **0.63** |
|  | **Demographic, Social** | **3** | **41.98** | **2.19** | **-17.47** | **0.14** | | **0.77** |
|  | **Demographic, Experience** | **3** | **42.31** | **2.53** | **-17.64** | **0.12** | | **0.89** |
|  | **Social, Experience** | **3** | **43.54** | **3.76** | **-18.25** | **0.06** | | **0.95** |
|  | Demographic, Social, Experience | 4 | 44.74 | 4.96 | -17.46 | 0.04 | | 0.99 |
|  | *null* | 1 | 47.60 | 7.82 | -22.74 | 0.01 | | 1.00 |
|  | Experience | 2 | 49.37 | 9.59 | -22.49 | 0.00 | | 1.00 |

| *16 days* | *Model names* | *K* | *AICc* | *ΔAICc* | *Log-likelihood* | *Weight* | *Cumulative weight* | |
| --- | --- | --- | --- | --- | --- | --- | --- | --- |
|  | **Demographic** | **2** | **39.78** | **0.00** | **-17.64** | **0.40** | | **0.40** |
|  | **Social** | **2** | **40.99** | **1.21** | **-18.25** | **0.22** | | **0.63** |
|  | **Demographic, Social** | **3** | **41.84** | **2.05** | **-17.40** | **0.14** | | **0.77** |
|  | **Demographic, Experience** | **3** | **42.31** | **2.53** | **-17.64** | **0.11** | | **0.88** |
|  | **Social, Experience** | **3** | **43.37** | **3.58** | **-18.16** | **0.07** | | **0.95** |
|  | Demographic, Social, Experience | 4 | 44.61 | 4.82 | -17.39 | 0.04 | | 0.99 |
|  | *null* | 1 | 47.60 | 7.82 | -22.74 | 0.01 | | 1.00 |
|  | Experience | 2 | 49.37 | 9.59 | -22.49 | 0.00 | | 1.00 |

| *18 days* | *Model names* | *K* | *AICc* | *ΔAICc* | *Log-likelihood* | *Weight* | *Cumulative weight* | |
| --- | --- | --- | --- | --- | --- | --- | --- | --- |
|  | **Demographic** | **2** | **39.78** | **0.00** | **-17.64** | **0.40** | | **0.40** |
|  | **Social** | **2** | **40.90** | **1.12** | **-18.20** | **0.23** | | **0.63** |
|  | **Demographic, Social** | **3** | **41.82** | **2.03** | **-17.39** | **0.14** | | **0.77** |
|  | **Demographic, Experience** | **3** | **42.31** | **2.53** | **-17.64** | **0.11** | | **0.88** |
|  | **Social, Experience** | **3** | **43.28** | **3.50** | **-18.12** | **0.07** | | **0.95** |
|  | Demographic, Social, Experience | 4 | 44.59 | 4.80 | -17.38 | 0.04 | | 0.99 |
|  | *null* | 1 | 47.60 | 7.82 | -22.74 | 0.01 | | 1.00 |
|  | Experience | 2 | 49.37 | 9.59 | -22.49 | 0.00 | | 1.00 |

| *21 days* | *Model names* | *K* | *AICc* | *ΔAICc* | *Log-likelihood* | *Weight* | *Cumulative weight* | |
| --- | --- | --- | --- | --- | --- | --- | --- | --- |
|  | **Demographic** | **2** | **39.78** | **0.00** | **-17.64** | **0.41** | | **0.41** |
|  | **Social** | **2** | **40.96** | **1.17** | **-18.23** | **0.23** | | **0.64** |
|  | **Demographic, Social** | **3** | **42.02** | **2.23** | **-17.49** | **0.13** | | **0.77** |
|  | **Demographic, Experience** | **3** | **42.31** | **2.53** | **-17.64** | **0.12** | | **0.89** |
|  | **Social, Experience** | **3** | **43.38** | **3.60** | **-18.17** | **0.07** | | **0.95** |
|  | Demographic, Social, Experience | 4 | 44.79 | 5.00 | -17.48 | 0.03 | | 0.99 |
|  | *null* | 1 | 47.60 | 7.82 | -22.74 | 0.01 | | 1.00 |
|  | Experience | 2 | 49.37 | 9.59 | -22.49 | 0.00 | | 1.00 |

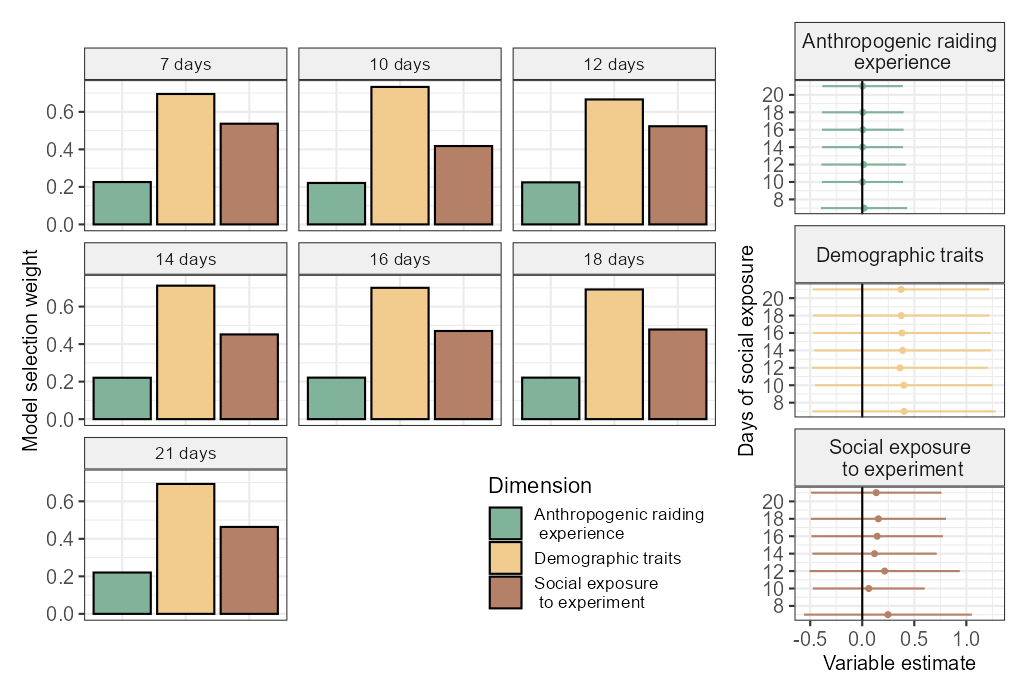

**Figure S12**. Comparison of models investigating the effects of the anthropogenic raiding experience, demographic, and social exposure to the box dimensions on good technical innovation scores, where the social exposure to the experiment varies. In the original analysis, we used a 14-day period (in green) of social exposure to the experiment. Here, we tested the sensitivity of this period by varying the number of days from 7 to 21.

*11.3 ‘good IT’ model: IT threshold*

**Table S7**. Comparison of models investigating the effects of the anthropogenic raiding experience, demographic, and social exposure to the experiment dimensions on good technical innovation scores (*IT*) with varying cut-offs. We used a binary version of good/bad technical innovation, based on whether the 95% confidence interval of the estimated *IT* parameter crossed the 0.5 line, the selection table of which is indicated in green. We tested the sensitivity of this cut-off here by varying this confidence interval cut-off from 0.3 to 0.7. *K* represents the number of estimated parameters for each model, *AICc* the Akaike Information Criterion corrected for small samples, *∆AIC* is obtained by comparing to the most parsimonious model (first row), log-likelihood is the probability of the observed data, *weight* corresponds to the Akaike weights (exp(-0.5**AICc*)), and their cumulative weight.

| *IT < 0.3* | *Model names* | *K* | *AICc* | *ΔAICc* | *Log-likelihood* | *Weight* | *Cumulative weight* | |
| --- | --- | --- | --- | --- | --- | --- | --- | --- |
|  | **Social, Experience** | **3** | **25.63** | **0.00** | **-9.29** | **0.26** | | **0.26** |
|  | **Social** | **2** | **25.77** | **0.14** | **-10.63** | **0.25** | | **0.51** |
|  | **Demographic** | **2** | **26.15** | **0.52** | **-10.83** | **0.20** | | **0.71** |
|  | **Demographic, Experience** | **3** | **27.22** | **1.59** | **-10.09** | **0.12** | | **0.83** |
|  | **Demographic, Social** | **3** | **27.57** | **1.94** | **-10.26** | **0.10** | | **0.93** |
|  | **Demographic, Social, Experience** | **4** | **28.37** | **2.74** | **-9.28** | **0.07** | | **1.00** |
|  | Experience | 2 | 42.00 | 16.37 | -18.80 | 0.00 | | 1.00 |
|  | *null* | 1 | 42.61 | 16.98 | -20.24 | 0.00 | | 1.00 |

| *IT < 0.4* | *Model names* | *K* | *AICc* | *ΔAICc* | *Log-likelihood* | *Weight* | *Cumulative weight* | |
| --- | --- | --- | --- | --- | --- | --- | --- | --- |
|  | **Demographic** | **2** | **33.13** | **0.00** | **-14.32** | **0.28** | | **0.28** |
|  | **Social** | **2** | **33.25** | **0.12** | **-14.38** | **0.26** | | **0.54** |
|  | **Demographic, Social** | **3** | **33.90** | **0.77** | **-13.43** | **0.19** | | **0.72** |
|  | **Social, Experience** | **3** | **34.61** | **1.48** | **-13.78** | **0.13** | | **0.86** |
|  | **Demographic, Experience** | **3** | **35.45** | **2.31** | **-14.20** | **0.09** | | **0.94** |
|  | **Demographic, Social, Experience** | **4** | **36.32** | **3.19** | **-13.25** | **0.06** | | **1.00** |
|  | *null* | 1 | 46.38 | 13.25 | -22.13 | 0.00 | | 1.00 |
|  | Experience | 2 | 46.94 | 13.81 | -21.27 | 0.00 | | 1.00 |

| *IT < 0.5* | *Model names* | *K* | *AICc* | *ΔAICc* | *Log-likelihood* | *Weight* | *Cumulative weight* | |
| --- | --- | --- | --- | --- | --- | --- | --- | --- |
|  | **Demographic** | **2** | **39.78** | **0.00** | **-17.64** | **0.42** | | **0.42** |
|  | **Social** | **2** | **41.14** | **1.35** | **-18.32** | **0.21** | | **0.63** |
|  | **Demographic, Social** | **3** | **41.98** | **2.19** | **-17.47** | **0.14** | | **0.77** |
|  | **Demographic, Experience** | **3** | **42.31** | **2.53** | **-17.64** | **0.12** | | **0.89** |
|  | **Social, Experience** | **3** | **43.54** | **3.76** | **-18.25** | **0.06** | | **0.95** |
|  | Demographic, Social, Experience | 4 | 44.74 | 4.96 | -17.46 | 0.04 | | 0.99 |
|  | *null* | 1 | 47.60 | 7.82 | -22.74 | 0.01 | | 1.00 |
|  | Experience | 2 | 49.37 | 9.59 | -22.49 | 0.00 | | 1.00 |

| *IT < 0.6* | *Model names* | *K* | *AICc* | *ΔAICc* | *Log-likelihood* | *Weight* | *Cumulative weight* | |
| --- | --- | --- | --- | --- | --- | --- | --- | --- |
|  | **Social** | **2** | **39.50** | **0.00** | **-17.50** | **0.46** | | **0.46** |
|  | **Demographic** | **2** | **41.64** | **2.14** | **-18.57** | **0.16** | | **0.62** |
|  | **Demographic, Social** | **3** | **41.86** | **2.37** | **-17.41** | **0.14** | | **0.77** |
|  | **Social, Experience** | **3** | **41.98** | **2.48** | **-17.47** | **0.13** | | **0.90** |
|  | **Demographic, Experience** | **3** | **44.18** | **4.68** | **-18.57** | **0.04** | | **0.94** |
|  | **Demographic, Social, Experience** | **4** | **44.62** | **5.13** | **-17.40** | **0.04** | | **0.98** |
|  | *null* | 1 | 46.38 | 6.89 | -22.13 | 0.01 | | 0.99 |
|  | Experience | 2 | 48.38 | 8.89 | -21.99 | 0.01 | | 1.00 |
| *IT < 0.7* | *Model names* | *K* | *AICc* | *ΔAICc* | *Log-likelihood* | *Weight* | *Cumulative weight* | |
|  | **Social** | **2** | **37.16** | **0.00** | **-16.33** | **0.27** | | **0.27** |
|  | **Demographic** | **2** | **37.30** | **0.14** | **-16.40** | **0.25** | | **0.52** |
|  | ***null*** | **1** | **38.68** | **1.53** | **-18.28** | **0.13** | | **0.65** |
|  | **Social, Experience** | **3** | **39.18** | **2.03** | **-16.07** | **0.10** | | **0.75** |
|  | **Demographic, Experience** | **3** | **39.34** | **2.18** | **-16.15** | **0.09** | | **0.84** |
|  | **Demographic, Social** | **3** | **39.69** | **2.53** | **-16.32** | **0.08** | | **0.91** |
|  | **Experience** | **2** | **40.07** | **2.91** | **-17.84** | **0.06** | | **0.98** |
|  | Demographic, Social, Experience | 4 | 41.94 | 4.78 | -16.06 | 0.02 | | 1.00 |

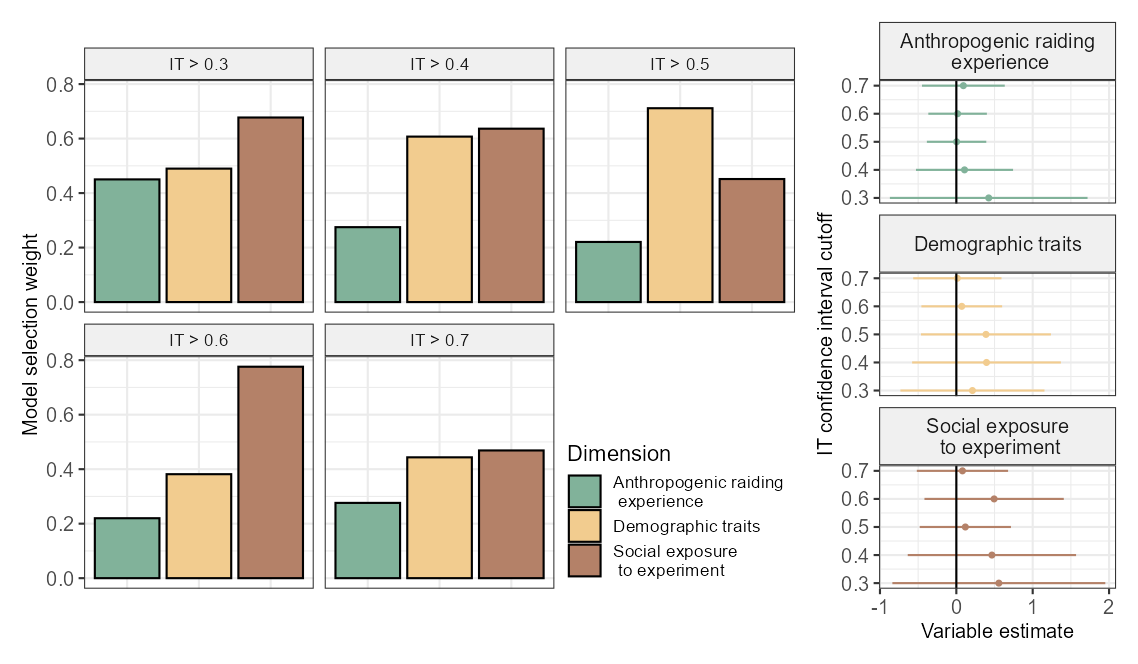
**Figure S13**. Comparison of models investigating the effects of the anthropogenic raiding experience, demographic, and social exposure to the experiment dimensions on good technical innovation scores (*IT*) with varying cut-offs. We used a binary version of good/bad technical innovation, based on whether the 95% confidence interval of the estimated *IT* parameter crossed the 0.5 line, the selection table of which is indicated in green. We tested the sensitivity of this cut-off here by varying this confidence interval cut-off from 0.3 to 0.7.

*12. Verification of models’ assumptions*

For the most complex models in our study, the *SS* model (see Section 2.5.3), ‘good *IT’* model (see Section 2.5.3), problem-solving success model (see Section 2.5.4), and anthropogenic feeding success model (see Section 2.5.4), we verified the model assumptions. We looked at the Variance Inflation Factors (VIFs, tables S8 – S11) and the histograms of residuals, the Q-Q plots, and the scatter plots of the fitted values vs. the residuals (figures S14 - S17). It highlighted no major issues.

*12.1 SS model assumption tests*

**Most complex model:**

*SS* ~ Anthropogenic raiding experience + Demographic traits + Social exposure to box

**Table S8.** Model assumptions check: ‘*SS*’ models | Variance Inflation Factor for the *SS* model using the “check_collinearity” of the *performance* package (Lüdecke et al. 2021). All VIFs are acceptable (<2).

| ***Term*** | ***VIF*** |
| --- | --- |
| Anthropogenic raiding experience | 1.307 |
| Demographic traits | 1.317 |
| Social exposure to experiment | 1.008 |

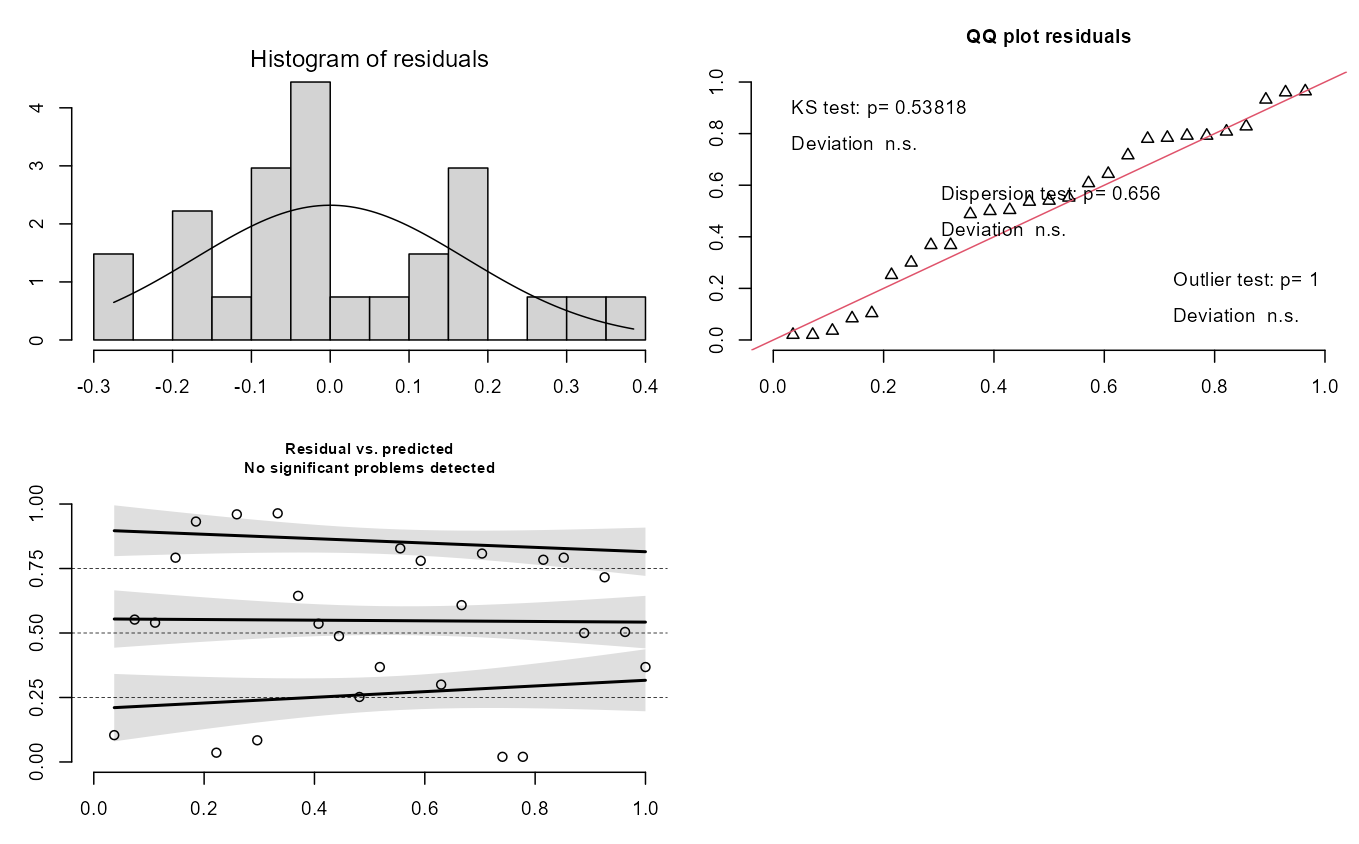

**Figure S14.** Model assumptions check: ‘*SS*’ models | Depicted are the histogram of residuals, the Q-Q plot (adapted for the modelled distribution; with test for deviation of the distribution (Kolmogorov-Smirnov, KS, outliers and overdispersion) and the scatter plot of the fitted values vs. the residuals. Plots were based on the *DHARMa* package (Hartig, 2022).

*12.2 good IT model assumption tests*

**Most complex model:**

good*IT* ~ Anthropogenic raiding experience + Demographic traits + Social exposure to box

**Table S9.** Model assumptions check: ‘good *IT*’ models | Variance Inflation Factor for the good *IT* model using the “check_collinearity” of the *performance* package (Lüdecke et al. 2021). All VIFs are acceptable (<2).

| ***Term*** | ***VIF*** |
| --- | --- |
| Anthropogenic raiding experience | 1.214 |
| Demographic traits | 1.205 |
| Social exposure to experiment | 1.013 |

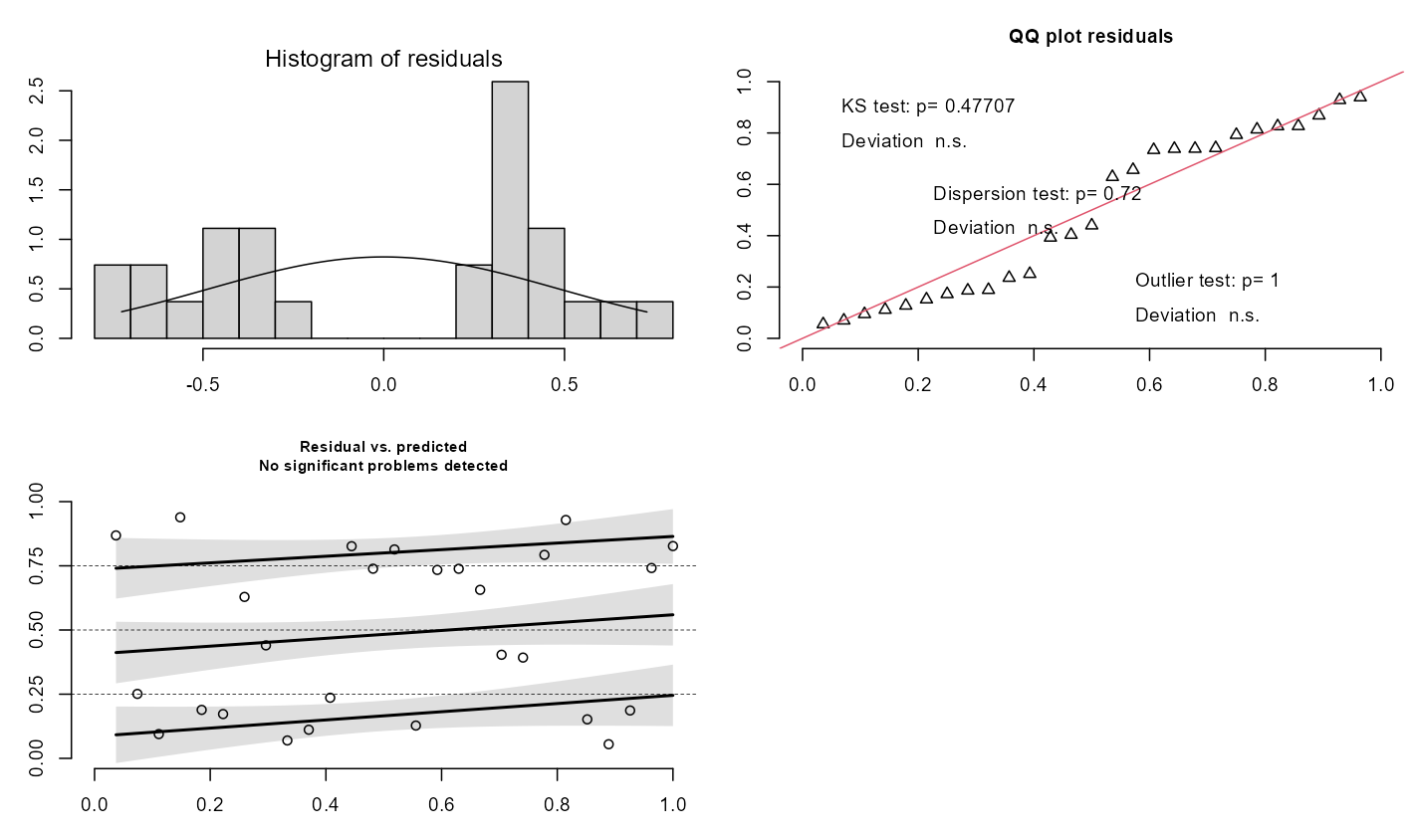

**Figure S15.** Model assumptions check: ‘good *IT*’ models | Depicted are the histogram of residuals, the Q-Q plot (adapted for the modelled distribution; with test for deviation of the distribution (Kolmogorov-Smirnov, KS, outliers and overdispersion) and the scatter plot of the fitted values vs. the residuals. Plots were based on the *DHARMa* package (Hartig, 2022).

*12.3 Problem solving success model*

**Most complex model:**

Problem solving success ~ *IT* + *SS^2^*

**Table S10.** Model assumptions check: problem-solving success model | Variance Inflation Factor for the success model using the “check_collinearity” of the *performance* package (Lüdecke et al. 2021). All VIFs are acceptable (<2).

| **Term** | **VIF** |
| --- | --- |
| *IT* | 1.112 |
| poly(*SS*, 2) | 1.112 |

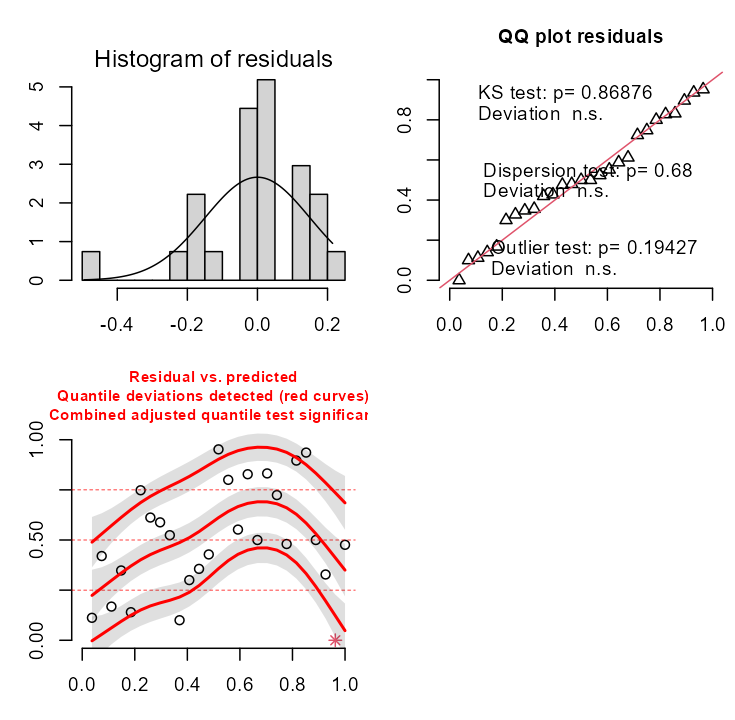

**Figure S16.** Model assumptions check: problem-solving success model | Depicted are the histogram of residuals, the Q-Q plot (adapted for the modelled distribution; with test for deviation of the distribution (Kolmogorov-Smirnov, KS, outliers and overdispersion) and the scatter plot of the fitted values vs. the residuals. Plots were based on the *DHARMa* package (Hartig, 2022). Although quantile deviations were detected the histogram and Q-Q plot did not reveal substantial departures from normality. Therefore, we consider the model assumptions to be sufficiently met for interpretation.

*12.4 Anthropogenic food success model*

**Most complex model:**

Anthropogenic food success ~ *IT* + *SS^2^*

**Table S11.** Model assumptions check: anthropogenic food success model | Variance Inflation Factor for the success model using the “check_collinearity” of the *performance* package (Lüdecke et al. 2021). All VIFs are acceptable (<2).

| **Term** | **VIF** |
| --- | --- |
| *IT* | 1.082 |
| poly(*SS*, 2) | 1.082 |

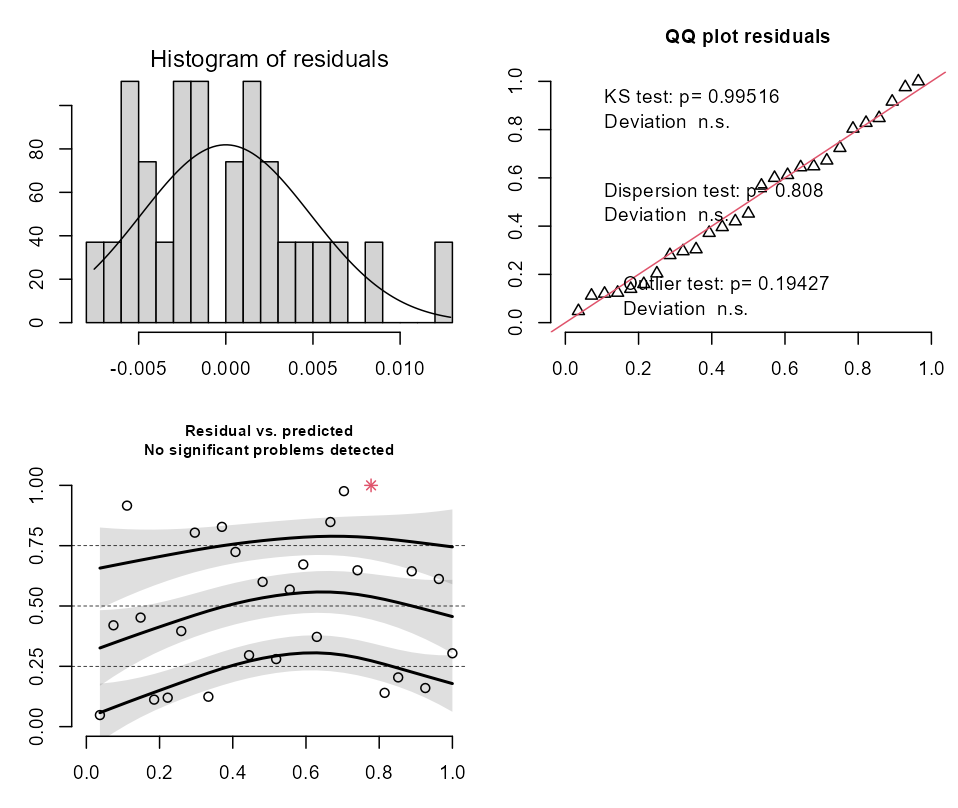

**Figure S17.** Model assumptions check: anthropogenic food success models | Depicted are the histogram of residuals, the Q-Q plot (adapted for the modelled distribution; with test for deviation of the distribution (Kolmogorov-Smirnov, KS, outliers and overdispersion) and the scatter plot of the fitted values vs. the residuals. Plots were based on the *DHARMa* package (Hartig, 2022).

*13. Behavioural flexibility syndrome model results*

**Table S12**. Results from the beta-regression models with logit error structures used to test for the presence of a behavioural flexibility syndrome.

| **Response** | **Predictor** | **Estimate** | **Std. Error** | **z value** | **p value** |
| --- | --- | --- | --- | --- | --- |
| *SS* | *IT* | -0.863 | 0.874 | -0.987 | 0.323 |
| *V* | *IT* | 1.688 | 0.746 | 2.263 | 0.024* |
| *V* | *SS* | -0.727 | 0.703 | -1.034 | 0.301 |
